# Mouse Fc-FcγRIV structure guides Fc engineering for cross-species FcγR recognition

**DOI:** 10.64898/2026.05.12.724433

**Authors:** Yakendra Bajgain, Mo Guo, Kelli M. Hager, Annalee W. Nguyen, Y. Jessie Zhang, Jennifer A. Maynard

## Abstract

Antibody-dependent cellular cytotoxicity (ADCC) is a major mechanism of action for many FDA-approved therapeutic antibodies that is driven by interactions between the antibody Fc and Fcγ receptors (FcγRs) on immune effector cells. Murine models used for preclinical antibody evaluation currently have limited predictive value for clinical ADCC performance due to interspecies differences in Fc-FcγR interactions. The molecular determinants governing Fc-FcγR engagement in mice remain poorly defined, complicating the interpretation of murine ADCC data and its clinical relevance. To address this, we present the high-resolution crystal structure of the receptor that regulates Fc-mediated cytotoxicity in mice, mouse FcγRIV, alone and in complex with mouse IgG2a Fc. This complex preserves key features of the human IgG1 Fc-human FcγRIIIa interface which mediates ADCC in humans including salt bridges, hydrogen bonds, and a proline sandwich. However, subtle variations in receptor orientation, Fc-FcγR electrostatics, and glycan positions reduce human IgG1 Fc- mouse FcγRIV binding affinity, resulting in species-restricted Fc-FcγR mediated immune responses. Modeling of human IgG1 Fc interactions with mouse FcγRIV predicted steric clashes, suggesting opportunities to modulate the interaction. One structure-guided substitution variant of human IgG1, Fc_humo_, maintains comparable human FcγRIIIa engagement with enhanced binding to and activation of mouse FcγRIV, relative to human IgG1 Fc. This study provides proof-of-concept for engineering human Fc domains for cross-species FcγR recognition and provides a strategic framework to improve the predictive power of *in vivo* preclinical models.

## INTRODUCTION

The vertebrate immune system relies on a coordinated network of cellular and molecular components to mount effective responses against invading pathogens and abnormal host cells.^1–3^ Within this intricate network, antibodies, especially those of the immunoglobulin G (IgG) isotype, play important roles in translating antigen recognition into effective immune engagement.^4^ In humans, four IgG subclasses (IgG1, IgG2, IgG3 and IgG4) interact with a family of activating and inhibitory Fc gamma receptors (FcγRs) on immune cells via their fragment crystallizable (Fc) region to trigger key effector responses that define the therapeutic efficacy of many antibodies.^5–9^

Among these receptors, human FcγRIIIa (hFcγRIIIa) is especially important due to its ability to trigger robust antibody-dependent cellular cytotoxicity (ADCC) by natural killer (NK) cells in response to human IgG1 Fc (hFc) presentation on opsonized aberrant cells.^8,10^ ADCC is a key killing mechanism of many FDA-approved therapeutic antibodies (e.g., trastuzumab, rituximab, and cetuximab).^8,10,11^ As the clinical application of antibodies has expanded, Fc engineering to modulate Fc-FcγR interactions for decreased (e.g., the LALAPG hFc variant) or increased activity (e.g., the GASDALIE and afucosylated hFc variants) has emerged as a key strategy to enhance therapeutic and safety profiles.^12–14^ A pivotal advance in Fc engineering was determination of the crystal structure of the hFc region in complex with hFcγRIIIa, which revealed the precise molecular interface governing hFc-hFcγRIIIa engagement and laid the foundation for subsequent structure-guided Fc optimization.^15^

Mouse models remain indispensable for preclinical evaluation of antibody efficacy and safety, yet their predictive value for clinical outcomes remains modest. A contributing factor is the differential interactions of human Fc domains with mouse versus human FcγRs.^16–19^ In mice, four IgG subclasses (IgG1, IgG2a, IgG2b, and IgG3) mediate immune responses by binding to FcγRs on immune cells, with IgG2a regarded as the most potent mediator of Fc-mediated cytotoxicity and functionally homologous to human IgG1.^4,20^ Like humans, mice express both activating (FcγRI, FcγRIII, FcγRIV) and inhibitory (FcγRIIb) FcγRs, but their expression patterns on immune cells differ between mice and humans, as does FcγR regulation of immune effector functions.^4^ While ADCC in humans is mediated by hFcγRIIIa on NK cells, it is regulated in mice primarily by the closest homolog, mouse FcγRIV (mFcγRIV) expressed on myeloid cells including macrophages, monocytes, and neutrophils.^20–22^ Even though murine NK cells express mFcγRIII, the dominant Fc-mediated responses are driven by mFcγRIV, which serves as a functional analog for hFcγRIIIa.^21,22^ Due to species-specific differences in Fc-FcγR interactions, human IgG1 antibodies exhibit reduced binding affinity and weaker activation of murine Fcγ receptors, including mFcγRIV, resulting in diminished antibody activity that lacks predictive utility when evaluated in immunocompetent mouse models.^16,20,23^

This translational gap highlights the need for improved mechanistic understanding of Fc-FcγR interactions in the murine system. Although mFcγRIV is central to immunity, the molecular basis for its engagement with mIgG2a-Fc remains undefined. Accordingly, we aimed to define the structural interactions between the mFcγRIV ectodomain and mIgG2a-Fc to provide the basis for murine Fc-mediated cytotoxicity and insights into hFc interactions with mFcγRIV in preclinical mouse models.^16^ These data were then used to establish a proof-of-concept for engineering cross-reactive Fc variants that could enable more accurate evaluation of ADCC-dependent therapeutic efficacy.

## RESULTS

### Human IgG1 Fc exhibits lower affinity for mFcγRIV compared to mouse IgG2a Fc

The Fc domain of murine IgG2a antibodies (hereafter called mFc) binds all mouse activating FcγRs, including mFcγRIV, with moderate-to-high affinity, while the hFc domain binds mFcγRIV with a weaker affinity.^24^ To precisely define these binding interactions, we expressed full-length antibodies with human 4D5 Fab arms (PDB: 1N8Z)^25^ that bind the HER2 cell surface receptor (**Fig S1A**) and mFc or hFc to generate 4D5 mFc and 4D5 hFc, respectively. For a quantitative comparison, we used biolayer interferometry (BLI) with antibodies captured by anti-CH1 FAB2G biosensor tips dipped into wells containing mFcγRIV at concentrations from 31–1000 nM. The resulting association and dissociation curves were fit to a 1:1 binding model to determine association (*k*_on_) and dissocation (*k*_off_) rate constants and the kinetic equilibrium disassociation constant (K_D_ = *k*_off_/*k*_on_) with steady-state analyses used to obtain the steady-state K_D,SS_. While 4D5 mFc exhibited a ∼25 nM affinity for mFcγRIV, 4D5 hFc had a >4-fold weaker equilibrium K_D_ of ∼115 nM for mFcγRIV (**Fig 1A,B, Table 1**). Since Fc-FcγR affinity correlates with Fc-mediated tumor cell killing, this K_D_ difference explains the poor cytotoxicity observed with human antibodies in mouse models^23,26^ and motivated us to define the structural basis for differential Fc engagement of mFcγRIV.

**Table 1.** Binding of 4D5 Fc variants to various receptors.

| | $k_{\text{on}} \pm \text{SD} (\times 10^5 \text{ M}^{-1} \text{ s}^{-1})$ | $k_{\text{off}} \pm \text{SD} (\times 10^{-2} \text{ s}^{-1})$ | $K_D \pm \text{SD} (\text{nM})$ | $K_{D,\text{ss}} \pm \text{SD} (\text{nM})$ |
| --- | --- | --- | --- | --- |
| **mouse FcγRIV** |  |  |  |  |
| 4D5 mFc | $7 \pm 2$ | $1.6 \pm 0.1$ | $25 \pm 5$ | $26 \pm 7$ |
| 4D5 hFc | $5 \pm 2$ | $5 \pm 3$ | $120 \pm 20$ | $110 \pm 10$ |
| 4D5 Fc<sub>humo</sub> | $5 \pm 2$ | $2.4 \pm 1.0$ | $55 \pm 9$ | $53 \pm 8$ |
| **human FcγRIIIa V158** |  |  |  |  |
| 4D5 hFc | $4.0 \pm 0.4$ | $4.9 \pm 0.8$ | $120 \pm 10$ | $120 \pm 8$ |
| 4D5 Fc<sub>humo</sub> | $4.4 \pm 0.7$ | $5.3 \pm 1.0$ | $120 \pm 20$ | $120 \pm 20$ |
| **human FcRn (pH 6.0)** |  |  |  |  |
| 4D5 hFc | $0.46 \pm 0.02$ | $2.0 \pm 0.1$ | $440 \pm 10$ | $120 \pm 30$ |
| 4D5 Fc<sub>humo</sub> | $0.44 \pm 0.04$ | $2.3 \pm 0.3$ | $520 \pm 60$ | $120 \pm 30$ |

Data shown here are average (n=3) ± SD.

**Figure 1.**
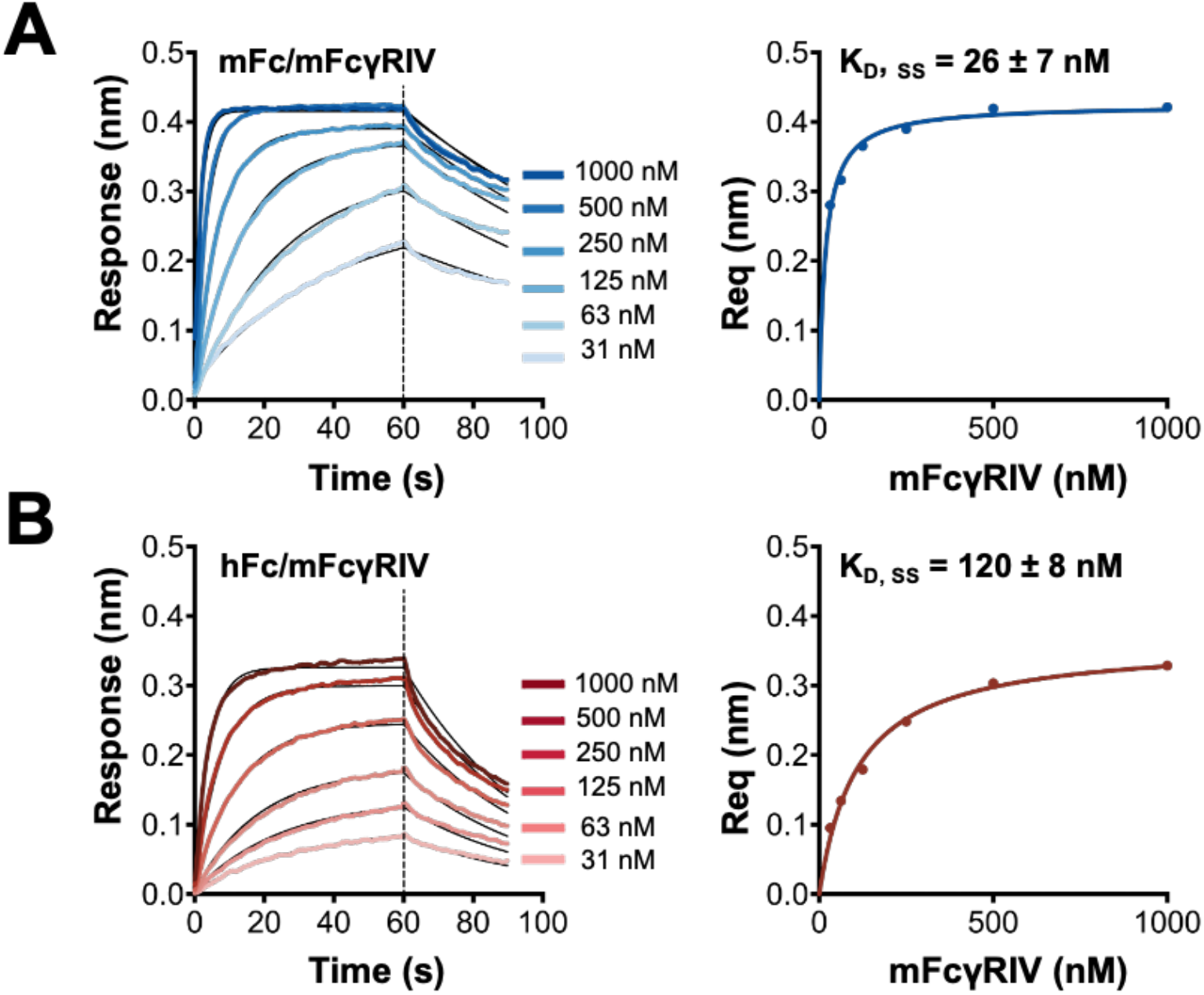
Human IgG1 Fc has a weaker binding to mouse FcγRIV compared to mouse IgG2a Fc. Biolayer interferometry (BLI) was used to quantify the binding of (***A***) mFc and (***B***) hFc with mFcγRIV using an OctRed96 instrument. Kinetic responses for the association and disassociation phases were fit to a 1:1 binding model, while the equilibrium responses were fit to a Langmuir isotherm. Data are representative of three independent experiments. K_D_ values from kinetic and equilibrium analyses are shown in Table 1.

### Structure of mouse FcγRIV reveals an Ig-like C2 domain organization

Mouse FcγRIV shares the highest sequence identity (∼65%) with hFcγRIIIa (**Fig. 2A**) among all human FcγRs, and both serve as the primary activating receptors mediating IgG-dependent effector cell cytotoxicity in their respective species.^27,28^ To characterize the mFcγRIV extracellular domain (Accession A0A0B4J1G0, residues 21-203), we determined its structure using protein expressed in Expi293 cells and purified via hexa-histidine affinity and sizing chromatographic steps. This resulted in a 36.5 kDa monodisperse, monomeric protein (**Fig. S1B**,**C**). Crystals formed in 0.1 M BIS-Tris (pH 5.5), 0.2 M MgCl_2_, and 28% polyethylene glycol 3350 at room temperature and diffracted to 2.8 Å resolution, with two molecules per asymmetric unit in space group P1 (**Table S1**). For molecular replacement, we used hFcγRIIIa (PDB: 1E4J, 65% amino acid identity)^15^ as a search model, excluding carbohydrates to avoid glycosylation bias. The electron density was continuous for all residues except for P32 in domain 1 (D1), resulting in a 2.8Å resolution model (**Fig. 2B**).

**Figure 2.**
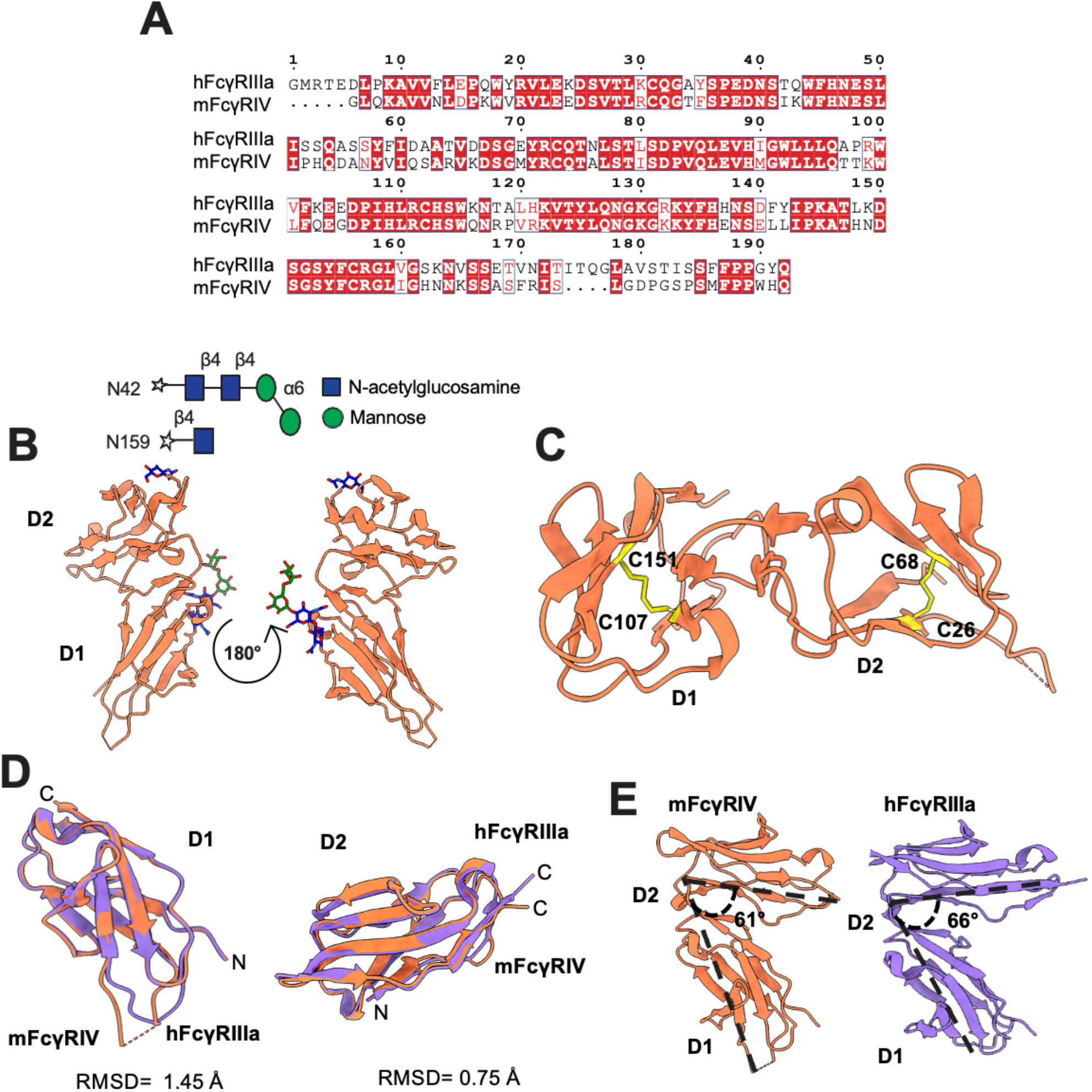
Structure of mouse FcγRIV reveals similarities to human FcγRIIIa. (***A***) The amino acid sequences of hFcγRIIIa V158 and mFcγRIV were aligned using Clustal Omega with default settings. Residues shown in red background are identical, and those in red text with white background are conserved residues. Overall, there is 65% identity between the two sequences. The alignment graphic was generated using ESPript. **(*B*)** Overview of the glycosylated mFcγRIV structure, shown in orange ribbon representation, with a 180° rotating view. Domains 1 and 2 are labeled as D1 and D2, respectively. N-linked glycans are depicted as sticks, with N-acetylglucosamine in blue and mannose in green. **(*C*)** Ribbon representation of the mFcγRIV backbone structure, highlighting the interstrand disulfide bond in yellow stick representation. **(*D*)** Structural superimposition of mFcγRIV (orange ribbon) and hFcγRIIIa (purple ribbon). The root-mean-square deviation (RMSD) of the main chain α-carbons for D1 and D2 are 1.45 Å and 0.75 Å, respectively and were calculated by superimposing the two structures based on Cα atoms using UCSF ChimeraX. **(*E*)** Comparison of the interdomain angles between domains 1 and 2 for mFcγRIV (orange ribbon) and hFcγRIIIa (light gray ribbon). The interdomain angle is indicated by a dashed black line and was measured in UCSF ChimeraX by defining planes for each domain using Cα atoms and calculating the angle between the planes.

Mouse FcγRIV adopts a characteristic fold with two immunoglobulin (Ig)-like C2 domains (D1 and D2), each forming a β-sheet sandwich stabilized by an intradomain disulfide bond (**Fig. 2C**). Its tertiary structure closely resembles hFcγRIIIa (PDB: 3SGJ) ^29^, which also features two extracellular Ig-like domains mediating IgG Fc interactions. While the individual domain structures of human and mouse are nearly identical (1.45 Å RMSD for D1, 0.75 Å RMSD for D2) (**Fig. 2D**), the interdomain orientation differs. Specifically, mFcγRIV adopts a 61° angle, slightly smaller than the 66° observed in hFcγRIIIa (**Fig. 2E**) which may influence receptor-ligand binding dynamics and functional interactions. Electron density maps revealed N-linked glycosylation sites at N42 and N159 (**Fig. 2B, Fig. S2**). The glycan at N159 included a single N-acetylglucosamine (GlcNAc), whereas N42 featured a more complex structure with two GlcNAc, one β-mannose (BMA), and one α-mannose (MAN) residue. For comparison, hFcγRIIIa contains five N-linked glycosylation sites in its extracellular domain (N38, N45, N74, N162, and N169), with glycan composition varying by cell type and expression systems.^30–32^ The aligned positions of mFcγRIV N42 and N159 correspond to hFcγRIIIa N45 and N162, respectively, suggesting conserved glycosylation sites across orthologs. Among hFcγRIIIa glycans, N162 has the greatest influence on hFc binding affinity because its glycan directly interacts with hFc N297 glycan and stabilizes the hFcγRIIIa binding interface to modulate ADCC.^32^ Oligomannose or minimally processed glycans at N162, like the ones present on NK cells, increase hFc-hFcγRIIIa binding affinity, whereas complex-type glycans tend to reduce binding affinity. In contrast, glycosylation at N45 can sterically hinder Fc engagement, while the remaining sites mainly contribute to receptor folding, stability, and surface expression.^30,31,33^ Overall, the mFcγRIV structure revealed a conserved Ig-like architecture with distinct interdomain geometry and simplified glycosylation relative to hFcγRIIIa. These features define mFcγRIV as the murine homolog for hFcγRIIIa and provide a structural basis for interpreting Fc-dependent effector functions of therapeutic antibodies in mouse models.

### Dimer opening enables mouse Fc to complex with mouse FcγRIV

Mouse IgG2a Fc shares ∼64% sequence identity with hFc and is the primary Fc isotype involved in Fc-mediated effector function in mice.^27^ The crystal structure of mFc was previously determined (PDB ID: 5VAA), with a T370K substitution that facilitates controlled Fab-arm exchange for production of murine bispecific antibodies.^34^ To better understand interactions between mFc and mFcγRIV, we obtained the crystal structure of unmodified mFc (UniProt Accession P01863, AA 98-330) alone and in complex with mFcγRIV. The mFc protein was expressed in Expi293 cells and purified using protein A and sizing chromatographic steps to yield a monodisperse protein with a 63.3 kDa molecular mass (**Fig. S1B**,**C**). Crystals grew in 0.1 M BIS-Tris (pH 6.0), 0.2 M LiSO4, and 20% PEG3350, with three molecules per asymmetric unit in space group C121 and diffracted to 3.2 Å resolution in a synchrotron (**Fig. 3A, Table S1**). The final model includes C_H_2 and C_H_3 domains (G237–S442), with clear electron density for N-glycans at position N297 in both chains (**Fig. 3A, Fig. S3A**,**B**) and disulfide bonds bridging residues C261–C321 in C_H_2 and residues C367–C425 in C_H_3 (**Fig. S3C**). Chain A features a bi-antennary complex-type oligosaccharide (GalGlcNAc_2_Man_3_GlcNAc_2_(Fuc)), while chain B carries the same bi-antennary complex-type oligosaccharide lacking one terminal galactose (GlcNAc_2_Man_3_GlcNAc_2_(Fuc); **Fig. 3A**).

**Figure 3.**
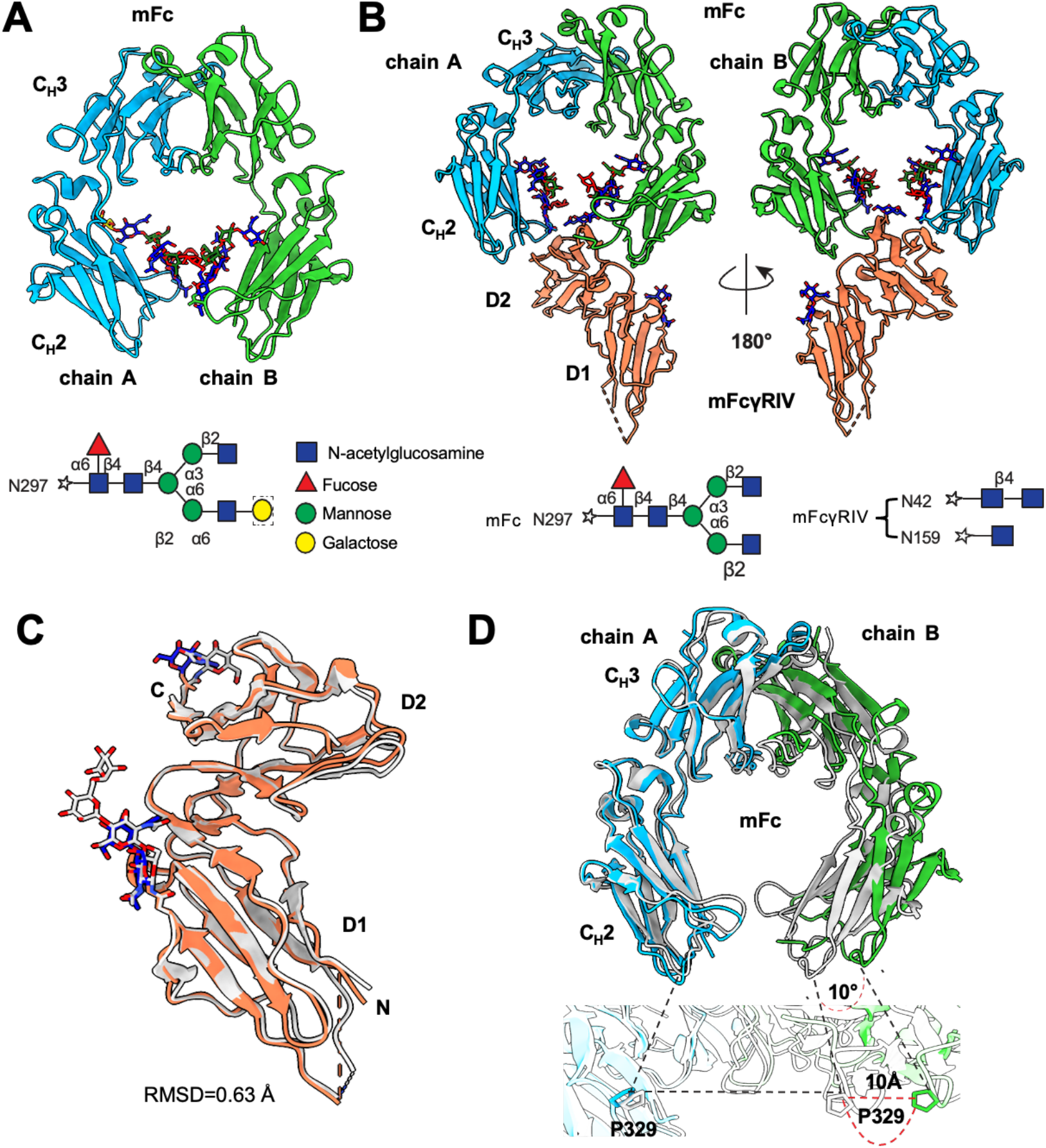
Structure of glycosylated mouse Fc- mouse FcγRIV complex shows conformational changes and Fc folds necessary for complex formation. (***A***) Standalone mFc homodimer structure. Monomer A is shown in light blue ribbon (labeled as chain A), and monomer B is shown in lime green ribbon (labeled as chain B). C_H_3 domain and C_H_2 domains are labeled as C_H_3 and C_H_2, respectively. N-linked glycans are depicted in stick representation, with N-acetylglucosamine in blue, mannose in green, fucose in red, and galactose in yellow. The box around galactose indicates presence of galactose on chain A but absence on chain B. (***B***) The mFc and mFcγRIV complex structure with a 180° rotating view. The mFc and mFcγRIV components are shown in the same colors as their standalone structures. N-linked glycans are depicted in stick representation, retaining the same color scheme as in Figure 2A. (***C***) Superimposition of the mFcγRIV complex structure and the standalone mFcγRIV structure. The mFcγRIV in the complex is shown in orange, while the standalone mFcγRIV is shown in light gray. The root-mean-square deviation (RMSD) of the main chain α-carbons is 0.63 Å. (***D***) *Top Panel*: Superimposition of the mFc complex structure and the standalone mFc structure. The Fc in the complex is shown using the same colors as in Figure 2A, while the standalone Fc is depicted in light gray. A black dashed curve highlights the shift of the C_H_2 domain in the complex structure relative to the standalone structure. *Bottom Panel*: Zoomed-in view of the C_H_2 region in contact with mFcγRIV. Residue P329 is shown in stick representation, and a black dashed line indicates the shifting distance of the C_H_2 domain between the complex and standalone structures.

To obtain the complex structure, purified mFc and mFcγRIV were combined and complexed species were isolated by size exclusion chromatography. The complex molecular weight was 96.2 kDa, consistent with a 1:1 mFc:mFcγRIV stoichiometry (**Fig. S1C**). Complex crystals were grown in 0.1 M BIS-Tris (pH 6), 0.1 M ammonium acetate and 20% polyethyleneglycol 8000, with one molecule per asymmetric unit in space group C2221 and diffracted to 3.16 Å (**Fig. 3B, Table S1**). Continuous electron density of the polypeptide backbone was observed for all residues except G28–P32 in D1 of mFcγRIV. The mFcγRIV structure remains nearly identical between its isolated and complexed forms, with a 0.63 Å RMSD for main-chain carbon atoms (**Fig. 3C**).^29^ The interdomain angle of mFcγRIV remains 61°, indicating that complex formation does not alter domain orientation. By contrast, the Fc undergoes asymmetric conformational changes upon complex formation. While preserving the overall fold, receptor binding pushes open the Fc lobes, increasing the P329–P329 distance at the C_H_2 domain tips by ∼10 Å to accommodate mFcγRIV (**Fig. 3D**). A similar asymmetric opening of the Fc C_H_2 lobes upon receptor engagement has been reported for the hFc-hFcγRI complex, where chain B undergoes a ∼9 Å outward displacement of P329 Cα relative to the apo structure (PDB: 4W4O).^35^ For the hFc-hFcγRIIIa complex, an asymmetric C_H_2 lobe opening has been reported, with conformational rearrangement of the lower hinge driving ∼7–8 Å outward displacement of the C_H_2 domain tips upon hFcγRIIIa engagement (PDB: 1E4K). ^15^

### The mFc-mFcγRIV complex is stabilized by electrostatic interactions and a proline-sandwich

The two mFc homodimer chains engage mFcγRIV asymmetrically, resulting in the commonly observed 1:1 binding stoichiometry for Fc-FcγR interaction.^35,36^ Mouse Fc chain A engagement with mFcγRIV is predominantly mediated by electrostatic interactions, while association with chain B is stabilized primarily through proline-sandwich stacking and hydrophobic contacts (**Fig. 4A**). Specifically, mFc chain A residues D269, D265, and D327 form salt bridges with mFcγRIV residues K128, K117, and H131, respectively. Residues S239 and S267 of mFc chain A additionally establish hydrogen bonds with mFcγRIV residues K117 and H131, while mFc chain A residues Y296 and N297 both form hydrogen bonds with mFcγRIV residue R152 (**Fig. 4B**). The interface between mFc chain B and mFcγRIV is stabilized predominantly by hydrophobic interactions and further reinforced by electrostatic contacts, including a salt bridge between mFcγRIV R113 and mFc D327 and hydrogen bonding between mFcγRIV N158 and mFc S239 (**Figs. 4C**). Residues W87 and W110 of mFcγRIV form a canonical proline-sandwich interaction with Fc residue P329, a defining structural feature of Fc-FcγR recognition. Additional hydrophobic residues G153, L154, and I155 pack against a hydrophobic LLGGP motif within mFc chain B, generating an extensive buried hydrophobic interface. In the complex, both mFc chains feature a bi-antennary complex-type oligosaccharide at N297 (GlcNAc_2_Man_3_GlcNAc_2_(Fuc)) (**Fig. 3B, Fig. S3D**). Mouse FcγRIV glycosylation again includes a single N-acetylglucosamine (GlcNAc) at residue N159 and a glycan structure at N42 with two N-acetylglucosamine residues (**Fig. 3B**). Notably, the β-mannose (BMA) and α-mannose (MAN) residue observed in the standalone receptor structure were not observed in the mFc-mFcγRIV complex, likely due to glycan heterogeneity or differences in electron density. Overall, this complex reveals that mFcγRIV recognition of mFc is achieved through asymmetric engagement and receptor-induced Fc opening, establishing the molecular basis for selective 1:1 Fc-FcγRIV binding underlying antibody-mediated immune regulation.

**Figure 4.**
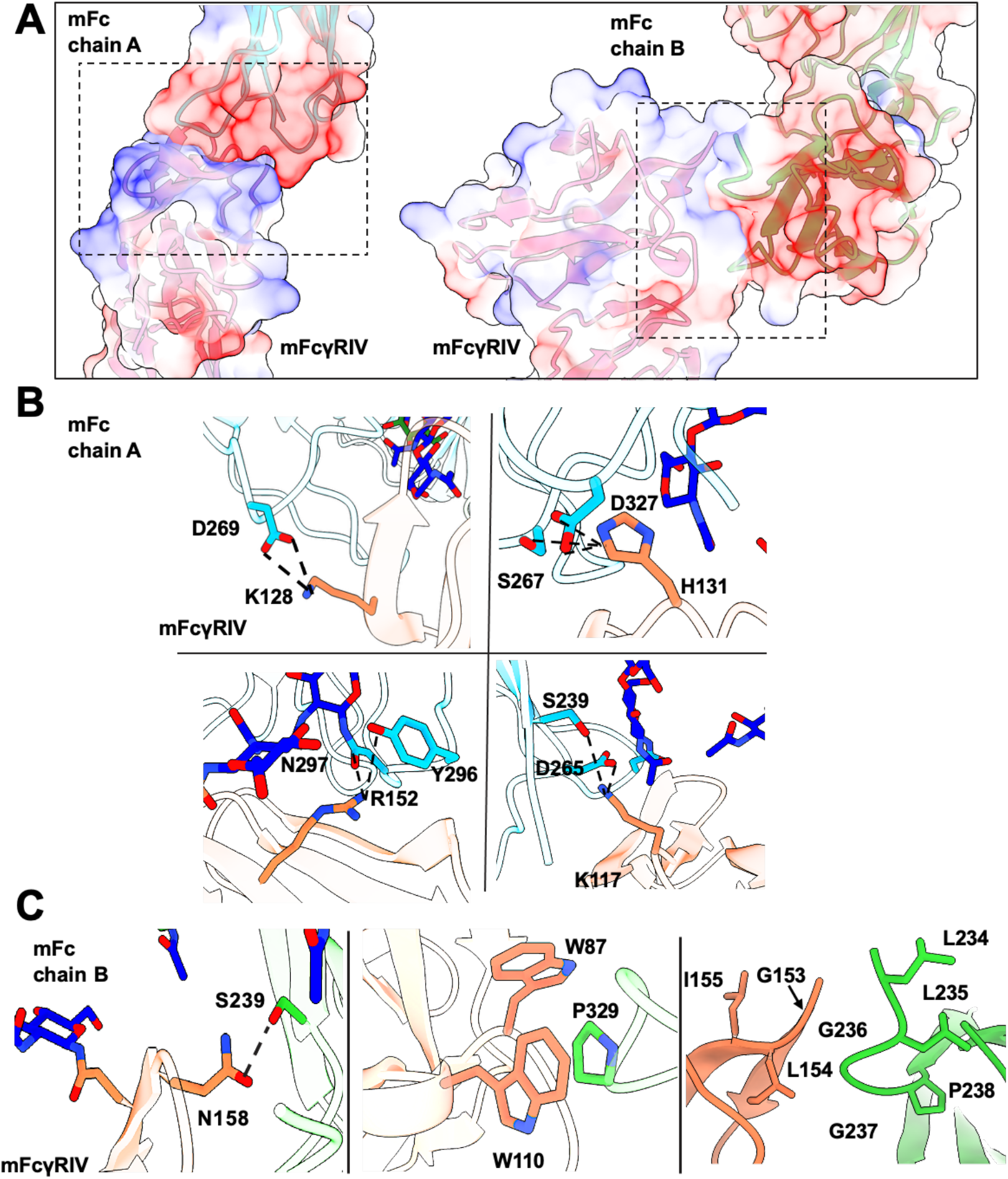
Mouse FcγRIV and mouse IgG2a Fc complex is stabilized by a proline sandwich and key electrostatic interactions. (***A***) Interface between mFcγRIV and mFc, shown with electrostatic surface representation. The charge distribution on the molecular surface is illustrated, with positively charged regions in blue, negatively charged regions in red, and neutral regions in white. This visualization highlights the electrostatic complementarity and potential interaction interfaces between mFcγRIV and the Fc. (***B***) Significant interactions within the interface between mFcγRIV and chain A. Mouse FcγRIV is shown in orange ribbon, and mFc Chain A is shown in light blue ribbon. Key residues (D269, K148, S267, D327, H151, N297, R172, Y296, K137, D265, and S239) are depicted as sticks. Black dashed lines indicate salt bridges or hydrogen bonds between interacting residues, and blue residue indicates N-acetylglucosamine. (***C***) Significant interactions within the interface between mFcγRIV and chain B. Mouse FcγRIV is shown in orange ribbon, and mFc chain B is shown in lime green ribbon. Key residues (N178, S239, R133, D327, W107, P329 and W130) are depicted as sticks. Black dashed lines indicate salt bridges or hydrogen bonds between interacting residues, and blue residue indicates N-acetylglucosamine.

### Distinct structural features modulate mFc-mFcγRIV versus hFc-hFcγRIIIa interactions

We compared our mFc-mFcγRIV structure with the published hFc-hFcγRIIIa (PDB: 3SGJ) complex using the Protein Interfaces, Surfaces, and Assemblies (PISA) program.^37^ A broadly similar asymmetric binding architecture is observed in both complexes, with differences in amino acid composition at the binding interfaces and N-linked glycan composition on the receptor. Specifically, chain A of the hFc-hFcγRIIIa interface has a comparable electrostatic interaction framework, with positively-charged hFcγRIIIa residues K131, H134, K120, and H135 interacting with negatively-charged hFc residues E269, D270, and D265 (**Fig. S4C**). Additional hydrogen bonds, including T122-N297, H134-S267, and K120-S239, further reinforce the human receptor-Fc interaction (**Fig. S4C**). The hFcγRIIIa-hFc chain B interface exhibits a similar proline-sandwich arrangement (W90-P329-W113) and hydrophobic packing between the receptor LVG residues and the Fc LLGGP motif as we observed in the murine interaction (**Fig. S4A, S4B, S4D**) but lacks an equivalent salt-bridge stabilization. Instead, the human interface is primarily supported by hydrogen bonds formed between receptor residues S160 and K114 with Fc residues E233 and N270 (**Fig. S4D**). Notably, the N159-linked glycan of mFcγRIV is positioned adjacent to the mFc chain A glycan but does not sterically interfere with receptor binding (**Fig. 3A**). In the hFc-hFcγRIIIa complex, the glycan at N162 of hFcγRIIIa makes extensive contacts with the hFc N297 glycan, forming a glycan-glycan interface that contributes to complex stability (**Table S2**).^38,39^ In contrast, the equivalent glycosylation site in mFcγRIV (N159) makes no significant interfacial contacts (<5 Å) with the mFc N297 glycan (**Table S2**). This comparison reveals that despite conservation of the overall complex architecture, differences in receptor electrostatics, glycosylation, and domain orientation generate receptor-specific Fc engagement interfaces between the two species.

### Predicted hFc binding to mFcγRIV reveals the molecular basis of differential affinity

To identify the structural basis for reduced binding of hFc to mFcγRIV, we superimposed the hFc structure (PDB: 3SGJ) with our mFc-mFcγRIV complex, focusing on chain A of the Fc region. This allowed us to directly compare the two Fc-FcγR interfaces while maintaining the experimentally determined receptor geometry. This comparison revealed three non-conserved residues, proximal to the mFcγRIV binding interface that differ in local side-chain polarity, electrostatic properties, and steric bulk (**Fig. 5A**). Residue 233 is glutamic acid in hFc, versus asparagine in mFc. Though E233 can form electrostatic interactions and hydrogen bonds with mFcγRIV residues R116 and H157, its longer side chain sterically hinders these residues from optimal interaction. Electron density for mFc N233 is absent, but structural alignment suggests that its hydroxyl group engages R116 and H157 with greater spatial complementarity (**Fig. 5B, C**), indicating that a polar residue at this position in hFc could relieve the steric strain while preserving electrostatic contacts. At position 268 of mFc, glutamic acid interacts with K128 of mFcγRIV but is replaced by histidine at the corresponding position in hFc (**Fig. 5D**), suggesting that restoring a negatively charged residue could recover this interaction. Finally, at position 327 in mFc, D327 forms extensive mFcγRIV contacts, with D327 in chain A forming a salt bridge with H131 and D327 in chain B forming a salt bridge with R113 and a hydrogen bond with Q111. The A327 in hFc is unable to form these critical contacts (**Fig. 5E**), indicating that introduction of a charged residue at position 327 in hFc may enhance hFc-mFcγRIV interactions. Together, these structural differences provide a molecular basis for the reduced hFc-mFcγRIV affinity and led us to speculate that targeted hFc substitutions could result in mFc-like engagement of mFcγRIV.

**Figure 5.**
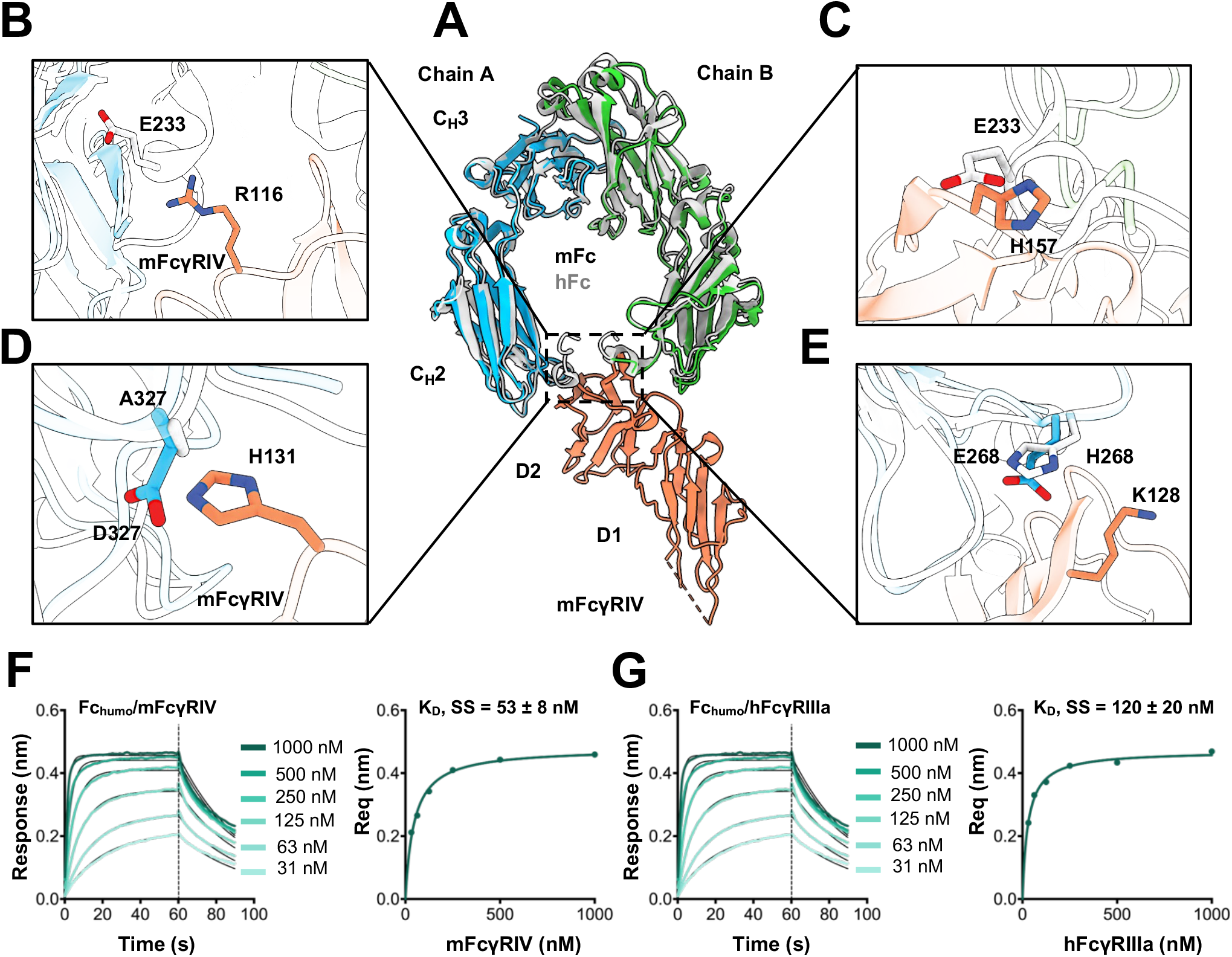
Structural analysis highlights key amino acid mismatches between human IgG1 Fc and mouse FcγRIV that weaken their interaction strength. ***(A*)** Superimposed structure of mFc and hFc interacting with mFcγRIV. Mouse FcγRIV is shown in orange ribbon, hFc chain A and chain B is shown in grey ribbons, and mFc chain A is shown in blue ribbon and chain B is shown in green ribbon. (***B-E***) The overall structure is accompanied by amino acid residues in hFc region (E233, H268, A327) within its binding interface with mFcγRIV that are different than mFc region and might explain the difference in their respective binding interactions. (***F-G***) BLI was used to quantify the binding of Fc_humo_ with (***F***) mFcγRIV and (***G***) hFcγRIIIa at various concentrations (31-1000nM) using an OctRed96 instrument. Kinetic responses for the association and disassociation phases were fit to a 1:1 binding model, while the equilibrium responses were fit to a Langmuir isotherm. Data are representative of three independent experiments. K_D_ values from kinetic and equilibrium analyses are shown in Table 1.

### E233D enhances hFc binding to mFcγRIV while retaining binding to hFcγRIIIa

To evaluate this hypothesis, we targeted hFc residues at positions 233, 268 and 327 for rational Fc engineering. Introduction of single alanine substitutions at positions 233 and 268 (**Fig. S5**), revealed that both hFc variants had reduced binding to mFcγRIV compared to unmodified hFc (**Fig. S6B**,**C**). We then introduced the corresponding mFc residues at each position individually (E233N, H268E, and A327D) to determine whether restoring an mFc-like chemical environment would improve binding (**Fig. S5**). However, these variants did not improve hFc-mFcγRIV binding based on ELISA results (**Fig. S6B**,**C**). We additionally substituted E233 with D (E233D, termed Fc_humo_ hereafter), which selectively enhanced binding to mFcγRIV relative to hFc without altering binding to hFcγRIIIa (**Fig. 5F, Fig. S6A**). Fc_humo_ expressed with similar yields (∼120 mg/L ExpiCHO culture) and was equally stable with unaltered thermal unfolding transition relative to hFc (**Fig. S7A**,**B**). This effect of the E233D substitution is consistent with its structural properties, as aspartic acid preserves the negative charge required for electrostatic complementarity with mFcγRIV residues while reducing side-chain length by one methylene group, relieving steric hindrance and restoring more optimal hydrogen-bonding geometry with the mFcγRIV interface residues.

To quantitatively assess binding of Fc_humo_ to the FcγRs mediating ADCC in humans and mice, we performed BLI with hFcγRIIIa and mFcγRIV. Whereas hFc bound mFcγRIV with a K_D,SS_ of ∼115 nM and mFc bound with ∼25 nM, Fc_humo_ exhibited an enhanced binding of K_D,SS_ of ∼55 nM (**Fig. S6D, Table 1**) compared to hFc (One-way ANOVA, Dunnett’s post hoc test, p=0.0006). In addition, Fc_humo_ maintained similar binding to both F158 and V158 hFcγRIIIa variants as hFc (**Table 1, Fig. 5G, Fig. S6E-G**) (Welch’s t-test, p= 0.807). The enhanced mFcγRIV binding and retained hFcγRIIIa engagement suggest that Fc_humo_ could serve as a cross-reactive platform in which the same molecule engages the relevant ADCC-mediating receptors in both species, making efficacy data generated in mouse models more informative about human biology.

### Fc_humo_ exhibits favorable binding to other human and mouse FcγRs

In humans, mutations designed to enhance binding to a specific FcγR often influence interactions with other FcγRs as well. This occurs because most FcγRs engage partially overlapping residues within the hinge and C_H_2 domains of the hFc, meaning that substitutions which stabilize a favorable Fc conformation typically increase affinity across multiple receptors simultaneously.^14,39^ For instance, the S239D/I332E mutations in Fc variants boost hFcγRIIIa affinity for improved NK-mediated ADCC but also enhances binding to hFcγRIIa on monocytes.^40,41^ To determine if the E233D substitution has similar effects, we assessed Fc_humo_ binding to other key human activating (hFcγRI, hFcγRIIa H131, and hFcγRIIa R131) and inhibitory (hFcγRIIb) receptors. We found that Fc_humo_ had similar binding affinities for all human activating and inhibitory FcγRs as hFc (**Figure S8, Table S3**). BLI analysis showed that hFc and Fc_humo_ bound hFcRn similarly at pH 6.0, with minimal binding at pH 7.4, indicating that the E233D mutation does not alter FcRn affinity under physiologically relevant conditions (**Fig. S7C–D, Table 1**). Consistent with this, both variants exhibited comparable serum half-lives in BALB/c mice (8.7±1.0 days for hFc versus 8.3±1.1 days for Fc_humo_ (**Fig. S7E**). This hFc-like hFcRn binding and similar serum half-life suggest that E233D substitution did not perturb the C_H_2-C_H_3 interface governing hFcRn engagement, indicating that its effects are confined to the classical FcγR interaction surface. Additionally, we determined the binding of Fc_humo_ with mouse activating and inhibitory FcγRs. Fc_humo_ exhibited ∼2-fold enhanced affinity for mFcγRI (**Fig. S9, Table S4**) (one-way ANOVA, Dunnett’s post hoc test, p= 0.0025), an activating receptor present on monocytes, macrophages, and dendritic cells that drives antibody-dependent phagocytosis and pro-inflammatory cytokine release in mice.^42,43^ We detected no noticeable binding of both hFc and Fc_humo_ to mFcγRIII (**Fig. S9, Table S4**). Binding to mFcγRIIB, the inhibitory receptor, was slightly lower than mFc but like how hFc bound. This selective enhancement of binding affinity for mFcγRI and mFcγRIV while retaining binding for hFcγRs suggests that substitution E233D exploits positional differences in how human versus mouse receptors contact the upper-lower hinge region of hFc C_H_2 domain.

### Fc_humo_ activates human FcγRIIIa and mouse FcγRIV similarly *in vitro*

ADCC is initiated by FcγR binding to clustered antibody Fc domains on target cells, leading to immune synapse formation, initiation of signaling and subsequent effector cell activation.^44–46^ Biochemical binding assays (ELISA, BLI) indicated that Fc_humo_ enhanced mFcγRIV binding while maintaining hFc-like hFcγRIIIa interactions. However, since biochemical Fc-FcγR interactions do not capture these multivalent interactions, we employed a cellular reporter assay in which activation of FcγR upon binding antibody-opsonized target cells induces luciferase expression (**Fig. 6A**).^47^ Using SKBR-3 target cells expressing high HER2 levels (∼10^6^ receptors/cell)^48^, serially diluted 4D5 antibodies with different Fc regions, and ADCC reporter cells, we observed similar robust hFcγRIIIa activation by hFc and Fc_humo_ (61±4 and 53±5 maximum fold-induction, respectively; **Fig. 6B, Table S5**), with both variants exhibiting similar EC_50_ values (**Table S5**). Using mFcγRIV reporter cells, mFc strongly activated the cells, while hFc showed low activation (22±5 and 2.0±0.5 maximum fold-change, respectively). Fc_humo_ induced intermediate activation (9±3), consistent with its intermediate binding affinity for mFcγRIV (**Fig. 6C, Table S5**). Both mFc and Fc_humo_ had comparable EC_50_ values, indicating similar potency of mFcγRIV engagement (**Table S5**). Since FcγR-driven NFAT signaling depends on receptor cross-linking on the effector cell surface, higher activation at saturation by mFc is consistent with more effective FcγRIV clustering. Notably, the relationship between Fc-FcγR affinity and effector cell activation was non-linear, as the approximately two-fold gain in binding affinity conferred by Fc_humo_ compared to hFc translated into a ∼4–5-fold enhancement in mFcγRIV-mediated cellular activation, reflecting the amplifying effect of avidity through multivalent receptor clustering at the target cell surface. Overall, Fc_humo_ preserves hFcγRIIIa effector function while enhancing mFcγRIV-mediated cellular activity compared to hFc, establishing it as a proof-of-concept dual-compatible Fc variant that may enable more faithful translation of ADCC-dependent antibody activities from mouse to human.

**Figure 6.**
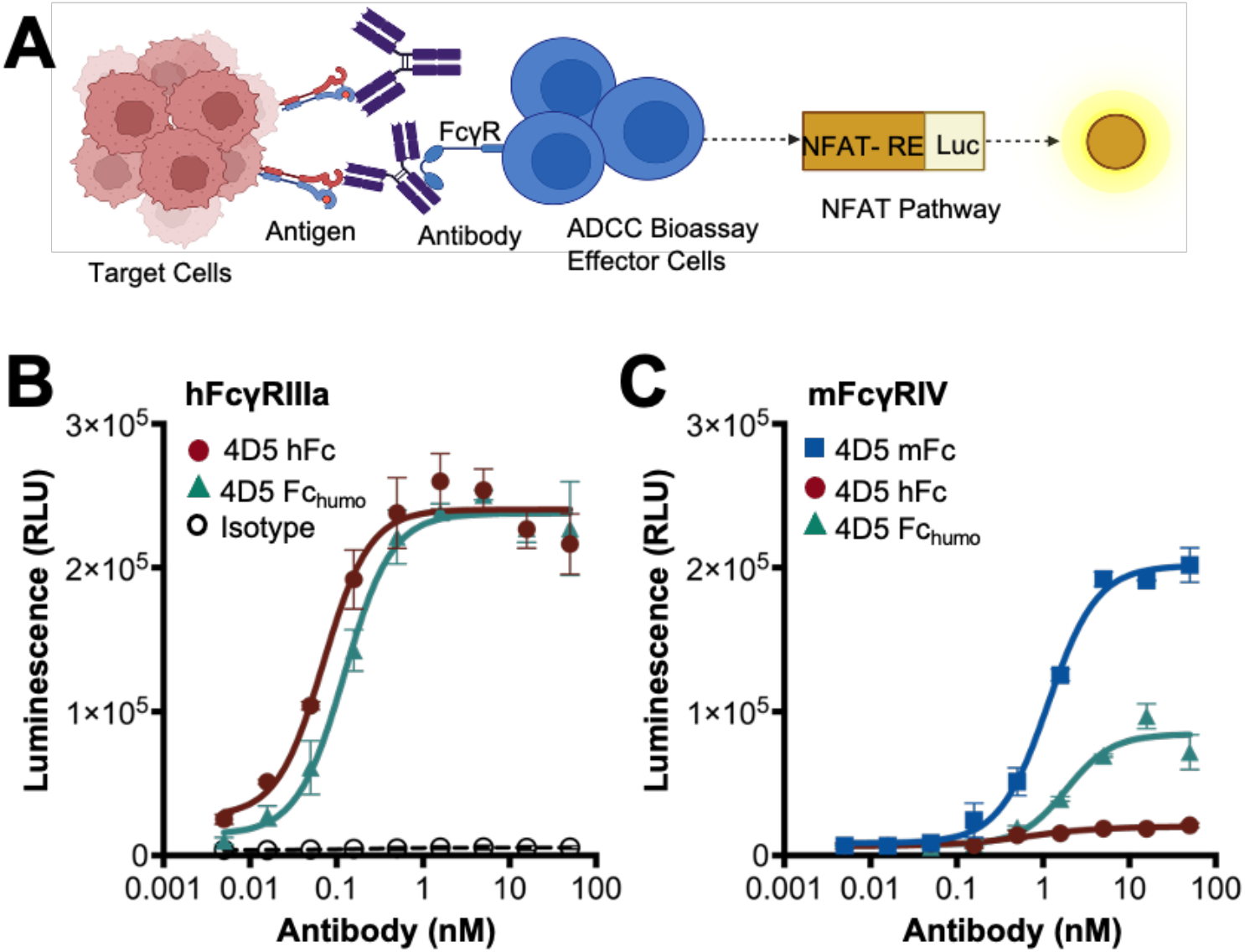
Fc_humo_ retains activation of human FcγRIIIa (V158) expressing effector cells and exhibits enhanced activation of mouse FcγRIV expressing effector cells. **(*A*)** Schematic representation of the ADCC assay. SKBR-3 cells expressing HER2 were used as target cells and incubated with effector cells at a 1:15 target:effector ratio for 18 h at 37 °C. Bio-Glo Reagent was added to measure luminescence. **(*B*)** Activation of hFcγRIIIa (V158) effector cells by increasing concentrations of anti-HER2 antibodies (4D5 variants). **(*C*)** Activation of mFcγRIV effector cells under the same conditions. Luminescence data were fitted to a four-parameter logistic (4PL) curve in GraphPad. Fold-induction relative to isotype or control antibodies are summarized in Table S5. Figures shown here are representative of three replicates.

## DISCUSSION

In this study, we present the high-resolution crystal structure of mFc in complex with the extracellular domain of mFcγRIV. In humans, the molecular basis of the hFc-hFcγRIIIa interaction has been described by high-resolution structures and validated by engineered hFc variants with enhanced or diminished receptor binding. The extracellular domain of mFcγRIV, the primary activating receptor for mIgG2a/2b mediated cytotoxicity, shares ∼65% sequence identity with that of hFcγRIIIa.^4,24^ Similarly, the mouse IgG2a Fc region shares ∼65% sequence identity with human IgG1 Fc, together positioning these Fc-FcγR pairs as functional orthologs that mediate Fc-mediated effector responses in each species. Despite this, the molecular basis for the mFc-mFcγRIV interaction has not been described, limiting our understanding of a key interaction mediating immune responses in mice. Our study provides a structural overview of this interaction, enables comparison with the well-characterized hFc-hFcγRIIIa complex and helps explain why many human antibody therapeutics exhibit reduced ADCC potency in mice despite strong activity in humans. Moreover, guided by the comparative structural analysis, we engineered Fc_humo_, which exhibits increased binding to mFcγRIV while retaining native affinity for hFcγRIIIa, serving as a proof-of-concept for structure-guided engineering of cross-species compatible Fc region for *in vivo* studies.

Comparison of the hFc-hFcγRIIIa and mFc-mFcγRIV complexes reveals a largely conserved overall architecture, with subtle structural differences contributing to species-specific Fc-FcγR binding. Compared to hFcγRIIIa, mFcγRIV adopts a smaller interdomain angle (61° versus 66°; **Fig. 2D**), which may influence its orientation toward the Fc homodimer and its engagement with the lower hinge and the CH2 region.^49^ Interestingly, the mFcγRIV interdomain angle does not change with complex formation, which contrasts with hFcγRIIIa, where the D1 and D2 interdomain angle increases as part of the structural rearrangement that occurs during the formation of the hFc-hFcγRIIIa complex.^15^ This distinction suggests a more rigid mode of mFc-recognition by mFcγRIV compared to an induced-fit-like mechanism observed in hFc-hFcγRIIIa complex. Like hFc-hFcγRIIIa complex, the mFc-mFcγRIV complex is also stabilized by an extensive network of salt bridges and hydrophobic interactions that, together with conserved features such as the proline-sandwich and LLGGP motif, result in their intermediate/high affinity interaction. However, the mFc-mFcγRIV chain B interface contains additional electrostatic stabilization through a salt bridge between R113 and D327, whereas the corresponding hFc-hFcγRIIIa chain is supported primarily by hydrogen bonding between interacting residues (**Fig. 3, Fig. S4**). Glycan architecture further differentiates the two interfaces. In mFcγRIV, the N159 glycan is positioned adjacent to the mFc chain A glycan in a manner that does not impose steric constraints with N297 glycan. In contrast, the N162 glycan of hFcγRIIIa is positioned proximal to the hFc N297 region and influences the local interaction environment through glycan-mediated contacts and modulation of Fc engagement geometry.^33,38^ This may affect Fc-FcγR binding within immune complexes and receptor clustering efficiency at the cell surface. FcγR signaling is highly dependent on nanoscale receptor clustering, which has been shown to enhance ITAM phosphorylation and downstream signaling.^49,50^ These structural differences in interface geometry and glycan positioning suggest a lower activation threshold for IgG-mediated Fc receptor crosslinking in mice for activation.

Antibody engineering approaches have been explored to enhance compatibility with murine systems for accurate translation of human effector mechanisms. Studies have shown that afucosylated hFc antibodies significantly enhance ADCC by anti-cancer antibodies through improved activation of mFcγRIV in luciferase-based ADCC assays.^28,51^ However, afucosylation of hFc also increases binding to hFcγRIIIa and therefore enhances ADCC in human systems as well, limiting its utility as a species-selective solution.^7,52^ Through our comparative structural analysis, we show that key residue mismatches in hFc (E233, H268, A327) in proximity with mFcγRIV residues introduce steric and electrostatic hindrances that alter its interaction with mFcγRIV. Guided by these insights, we explored the feasibility of engineering a human- and mouse-compatible Fc variant. Substitutions were selected to relieve predicted steric and electrostatic clashes at the hFc-mFcγRIV interface while preserving the native hFc fold and its interaction with hFcγRIIIa. Notably, these amino acid positions (E233, H268, A327) have previously been targeted to fine-tune FcγR interactions. E233D and H268D were identified, in combination with other mutations, to confer selective enhancement of hFcγRIIb binding over other activating hFcγRs.^53^ H268D was also incorporated into a Fc variant that preferentially activate hFcγRIIIa-mediated ADCC in the acidic tumor microenvironment.^54^ A327G selectively reduces hFc affinity for hFcγRIIIb and is also a part of complement silencing variants in clinical antibodies like Crovalimab^55,56^, which underscores their functional importance at the Fc-FcγR interface. Consistent with our structural predictions, Fc_humo_ enhanced binding to mFcγRIV by two-fold while retaining binding to hFcγRIIIa. Residue 233 lies immediately N-terminal to the lower hinge hFc residues L234 and L235, which are critical determinants of Fc binding to both human and mouse FcγRs. The bulkier glutamic acid at this position in hFc likely imposes steric strain that propagates into the lower hinge to subtly distort the geometry of L234 and L235 away from their optimal receptor-engaging conformation. Fc_humo_ preserves local electrostatic charge but shortens the side chain by one methylene group, which optimizes steric fit and local shape complementarity to enhance electrostatic interactions with key mFcγRIV residues (R116, H157) in the binding interface. Human IgG1 and mouse IgG2a are functional orthologs, and our structural analyses show that they engage their respective FcγRs through a conserved binding mechanism, consistent with the observation that LALAPG-mediated effector silencing is recapitulated across species.^57^ This suggests that cross-species differences in mFcγRIV engagement do not reflect a fundamentally altered binding mode, but rather the cumulative effect of a few divergent residues at a shared interface amenable to modulation.

Despite widespread use of mouse models for preclinical antibody evaluation, many human antibodies fail to elicit robust antibody-mediated killing of tumor cells in immunocompetent, syngeneic mice. This limitation arises from fundamental differences in FcγR composition, expression patterns, and binding properties between the two species.^19,20,58^ Human IgG1 binds mouse FcγRs with lower affinity than mouse IgG2a/2b, which may limit receptor clustering and reduce downstream signaling. For example, in one study, the anti-EpCAM antibody Adecatumumab showed potent ADCC with human effector cells but minimal activity with murine splenocytes *in vitro* and poor *in vivo* efficacy using a melanoma model. When its human IgG1 Fc was replaced with a murine IgG2a Fc, strong tumor-killing was observed and antibody treatment eliminated lung metastases in a syngeneic B16-EpCAM model.^23^ To address these translational challenges, the entire human FcγR repertoire was introduced into FcγR-deficient mice to restore human effector functions *in vivo*, either in addition to or instead of endogenous mouse receptors.^59,60^ More recently, knock-in models have replaced mouse FcγRs with their human counterparts under physiologic regulatory control to preserve their cell-specific expression and endogenous promoter regulation.^16,58^ In one such model, human FcγR knock-in mice more accurately recapitulated the clinical effects of glycoengineered anti-CD20 antibody Obinutuzumab compared to Rituximab, which was a distinction conventional mouse models failed to capture.^16^ However, engineering and maintaining humanized models is complex and resource-intensive, which limits their accessibility to many researchers. As a result, there remains a need for complementary approaches, such as *ex vivo* tonsil organoid systems or Fc-engineering strategies to accurately evaluate human antibodies in simpler mouse models.^17,26,61^ Fc_humo_ may address this need by enabling evaluation of Fc-dependent antibody mechanisms in conventional syngeneic mouse models. It confers improved mFcγRIV binding affinity and enhanced cellular activation *in vitro* while fully preserving hFcγRIIIa engagement, making it a promising tool for preclinical studies where drawing human clinically relevant conclusions from syngeneic mouse models has been confounded by poor hFc-mFcγR engagement.

Here, we determined the crystal structures of mFcγRIV alone and in complex the mFc and identified the molecular features driving this interaction. Comparison of this complex structure with its human homolog, hFc-hFcγRIIIa, revealed species-specific differences in interface geometry and glycan positioning. We exploited these differences to engineer Fc_humo_, a human Fc variant that retains hFcγRIIIa binding with enhanced mFcγRIV engagement. The limitations of this work include the use of static structural modeling, which does not capture conformational dynamics or glycan-dependent effects that influence Fc-FcγR interactions. In addition, our functional analyses of Fc_humo_ relied on FcγR reporter bioassays, which do not recapitulate NK cell or macrophage-mediated ADCC, as they have high expression of the cognate receptor but lack the simultaneous crosstalk among activating and inhibitory FcγRs present in physiological settings.^62^ To address this, future work will include *in vivo* validation in tumor models dependent on mFcγRIV-mediated effector function such as the B16F10-gp75 melanoma model.^21^ Together, this work defines the structural determinants of species-specific Fc-FcγR interactions and demonstrates how these insights can be leveraged to engineer Fc variants with cross-species receptor engagement for cytotoxic effects.

## Acknowledgements

We acknowledge the use of the CPRIT Advanced Protein Therapeutics core, RRID SCR_023740 for instrumentation and antibody expression support. We would like to thank Diamond Light Source for beamtime (proposal mx83239), and the staff of beamlines I24 for assistance with crystal testing and data collection.

## Funding and additional information

This work was funded by the Cancer Prevention and Research Institute of Texas (RP220587 to J.A.M.), the Welch Foundation (F-1767 to J.A.M.), the National Institute of General Medical Sciences (R35148356 to Y.Z), American Cancer Society Institutional Research Grant (IRG-21-135-01-IRG to A.W.N.).

## METHODS

### Antibody expression and purification

Antibodies were produced by transiently transfecting Abvec plasmids containing light chain and heavy chain of 4D5 (anti-HER2 antibody) variants into ExpiCHO cells following manufacturer’s standard titer protocol. Conditioned medium was centrifuged at 3,200 × g for 30 min. Antibodies were purified by protein A affinity chromatography using a 1-mL HiTrap Protein A HP column (Cytiva) on an ÄKTA FPLC system. The column was equilibrated in binding buffer (25 mM Tris, 25 mM NaCl, pH 7.4) and antibodies were eluted in elution buffer (100 mM sodium citrate, 50 mM NaCl, pH 3.0). Peak fractions were pooled, buffer-exchanged into phosphate-buffered saline (PBS), and stored at 4°C.

### Human and mouse Fc and FcγRs expression and purification

Abvec plasmids containing mouse IgG2a Fc (Accession P01863) and human IgG1 Fc (Accession P01857) were expressed in ExpiHEK cells and purified using protein A column as described above. Supernatants were harvested after 7 days by centrifugation at 3,200 × g for 30 min. C-terminal Hexahistidine-tagged human FcγRs (hFcγRI, Accession P12314; hFcγRIIb, Accession P31994; hFcγRIIa H131R, Accession P12318; and hFcγRIIIa V158F, Accession P08637) and mouse FcγRs (mFcγRI, Accession P26151; mFcγRIIb, Accession P08101; mFcγRIII, Accession P08508; and mFcγRIV, Accession A0A0B4J1G0) were expressed in ExpiHEK cells as described above and purified by immobilized metal affinity chromatography (IMAC) using Ni-NTA agarose (Qiagen, catalog no. 30210). UniProt Accession numbers for all receptors are listed in the Key Resources Table. Columns were washed with wash buffer (50 mM NaH_2_PO_4_, 300 mM NaCl, 20 mM imidazole, pH 8.0) and eluted with elution buffer (50 mM NaH_2_PO_4_, 300 mM NaCl, 250 mM imidazole, pH 8.0). Protein-containing eluted fractions were pooled, buffer-exchanged into PBS, and stored at 4°C.

### Protein purity and homogeneity assessment

The purity and homogeneity of all purified antibodies, Fc fragments, and FcγRs were evaluated by SDS-PAGE and size-exclusion chromatography (SEC). Proteins (3 μg per lane) were loaded on 4-12% Bis-Tris gels under reducing (5% v/v β-mercaptoethanol) and non-reducing conditions and visualized by Coomassie staining. For SEC, proteins were injected onto a Superdex 200 Increase 10/300 GL column (Cytiva) equilibrated in PBS on an ÄKTA FPLC system. Elution was monitored by absorbance at 280 nm.

### Fc: FcγR complex generation

SEC purified mouse IgG2a Fc and mouse FcγRIV were mixed in a 1:1 molar ratio and incubated overnight at 4°C to allow the mIgG2a-Fc: mFcγRIV complex to form. The molecular weight, purity, and homogeneity of purified proteins as well as the complex were verified protein gel and SEC as described above.

### Crystallization

Sitting-drop vapor diffusion method was employed to obtain diffraction-quality crystals. For crystallization of mouse FcγRIV, 1 μL of protein at 8.5 mg/ml and reservoir solution containing 0.1 M BIS-Tris (pH 5.5), 0.2 M Magnesium Chloride and 28% polyethyleneglycol 3350 were mixed at 1:1 ratio in the HR3-159 Cryschem Plate (Hampton Research) at 22°C. For crystallization IgG2a-Fc, 1 μL of protein at 8 mg/ml and reservoir solution containing 0.1M Bis-Tris (pH 6.0), 0.2 M Lithium Sulfate and 20% PEG3350 were mixed at 1:2 ratio in the HR3-159 Cryschem Plate (Hampton Research) at 22°C. For crystallization of mouse Fc-FcγRIV complex, 1 μL of protein at 7.5 mg/ml and reservoir solution containing 0.1 M BIS-Tris (pH 6.0), 0.1 M Ammonium Acetate and 20% polyethyleneglycol 8000 were mixed at 1:1 ratio in the HR3-159 Cryschem Plate (Hampton Research) at 22°C. The final crystals for three constructs were obtained after 5 –7 days. Individual crystals were flash-frozen directly in liquid nitrogen after brief incubation with a reservoir solution supplemented with 30% (v/v) glycerol.

### Data collection and structure determination

X-ray diffraction data were collected at I24 beamline in Diamond Light Source (Oxfordshire, UK). FcγRIV and Fc data were collected using CdTe Eiger2 9 M detector at a wavelength of 0.6199 Å and Fc-FcγRIV complex data was collected using Pilatus 6M at a wavelength 0.9999 Å. All datasets were processed using XIA2-DIALS pipeline which is integrated in Diamond ISPyB auto processing platform. X-ray diffraction patterns were processed to 3.21 Å (FcγRIV), 2.81 Å (Fc), and 3.16 Å (Fc-FcγRIV complex) respectively. In Phenix software, phases of FcγRIV and Fc were obtained by molecular replacement using human FcγRIII-B (PDB: 1E4J) and mouse IgG2a-Fc (PDB: 5VAA) as search model. Phase of Fc-FcγRIV complex was obtained using previously obtained standalone structure of Fc and FcγRIV as search model. Phase information was derived from the protein cores, excluding carbohydrates, and the resulting electron density maps revealed clearly distinguishable density at the expected locations of the glycosylation sites. The saccharide chains were manually built in Coot. All structures were iteratively built using Coot and Phenix refinement package. The quality of the finalized crystal structures was evaluated by MolProbity. The final statistics for data collection and structural determination are shown in Supplementary Table 1.

### ELISA Binding Assay

96-well high-binding plates were coated overnight at 4°C with FcγRs at 2 μg/mL in PBS. Wells were blocked for 1 h at room temperature with 2% BSA in PBST (PBS + 0.05% Tween-20), then washed three times with PBST. Serially diluted 4D5 antibodies with different Fc variants were incubated for 1 h at room temperature. After three washes, bound antibodies were detected with goat F(ab’)_2_ anti-human kappa-HRP (1:1,000) for 1 h at room temperature. After three additional washes with PBST, 50 μL TMB substrate was added per well and the reaction was quenched with 50 μL of 1 N HCl. Absorbance at 450 nm was measured and data were fitted to a four-parameter logistic (4PL) model in GraphPad Prism.

### Biolayer interferometry (BLI)

Fc-FcγR binding kinetics were measured by BLI using FAB2G biosensor tips (Sartorius). Tips were hydrated in kinetic buffer (0.1% BSA in PBST) for 15 min and loaded with 4D5 antibody variants at 10 μg/mL. After a baseline step in kinetic buffer, association was measured by dipping sensors into serially diluted FcγRs, followed by dissociation in kinetic buffer for. Kinetic rate constants (*k*_*on*_, *k*_*off*_) were derived by fitting a 1:1 Langmuir model to the full association phase and the first 5 sec of dissociation. Equilibrium dissociation constants (*K*_*D*_) were determined from the Langmuir isotherm: *R*_*eq*_ *= R*_*max*_ *× C / (K*_*D*_ *+ C)*, where *R*_*eq*_ is the steady-state response at analyte concentration *C* and *R*_*max*_ is the maximum binding response.

### *In vivo* half-life determination

BALB/c mice (n = 5 per group) were injected intraperitoneally with 2 mg/kg of 4D5 antibody containing human IgG1 Fc, Fc_humo_, or mouse IgG2a Fc. Blood was collected from the tail or submandibular vein on days 0, 1, 4, 7, 11, 14, 17, 20, 23, and 26. A terminal cardiac puncture was performed on day 26. Three mice in the mFc group were euthanized on day 17 per predetermined humane endpoints due to tail necrosis. Whole blood was centrifuged at 2,000 × g for 10 min at room temperature, and serum was stored at -80°C until analysis. Serum antibody concentrations were quantified using an indirect ELISA as previously described. Briefly, 96-well plates were coated with 0.5 μg/mL HER2-Fc in PBS. After blocking, mouse serum samples were initially diluted 1:100 in blocking buffer and subjected to three 1:3 serial dilutions. All samples were analyzed in duplicate. Bound antibodies were detected using goat anti-human kappa-HRP diluted 1:2000. A standard curve of 4D5 with human IgG1 Fc was included on each plate to determine antibody concentrations in serum samples. Serum antibody concentrations were calculated using a four-parameter logistic fit of the standard curve generated with Microsoft Excel Solver. Individual mouse antibody concentrations were plotted as a function of time (days) and fit to a single-phase exponential decay model. The slope constant (*K*) was obtained from the fitted curve, and half-life (*t*_*½*_) was calculated using the equation *t*_*½*_ *= ln(0*.*5)/K*. The mean half-life for each group was calculated.

### ADCC reporter Bioassay

FcγR-mediated ADCC activity was measured using the ADCC Reporter Bioassay Propagation Kit (Promega), which employs Jurkat cells stably expressing human FcγRIIIa V158 or mouse FcγRIV coupled to NFAT-driven luciferase reporter. High-HER2 expressing SKBR-3 target cells were seeded at 10,000 cells per well in growth medium (100 μL DMEM supplemented with 10% FBS) in white opaque 96-well plates and allowed to adhere overnight at 37°C, 5% CO_2_. On the day of the assay, 95 μL of growth medium was aspirated and replaced with 25 μL of assay buffer per well. Assay buffer consisted of RPMI 1640 supplemented with 0.5% low-IgG FBS for the hFcγRIIIa assay or 4% low-IgG FBS for the mFcγRIV assay. Serially diluted antibody variants (25 μL) were added in duplicate, followed by 150,000 effector cells in 25 μL assay buffer (15:1 effector:target ratio). Plates were incubated for 16 h at 37°C, 5% CO_2_. Wells containing target and effector cells without antibody served as the no-antibody control, and wells containing assay buffer only served as background. After equilibration to room temperature for 15 min, 75 μL of Bio-Glo Luciferase Assay Reagent (Promega) was added per well and luminescence was measured after 15 min. Fold induction was calculated as *(RLU*_*sample*_ *– RLU*_*no-antibody*_*) / (RLU*_*no-antibody*_ *-RLU*_*background*_) and data were fitted to a 4PL model in GraphPad Prism v10.3.1.

### Statistics

All statistical analyses were performed in GraphPad Prism v11.0.1 (99). Data are presented as mean ± SD unless otherwise indicated. Comparisons between two groups were performed using Welch’s t-test. For comparisons among three groups, ordinary one-way analysis of variance (ANOVA), Dunnett’s post hoc test to compare each group to a control group. A p-value <0.05 was considered statistically significant. Slope constants for *in vivo* half-life were compared using one-way ANOVA followed by Tukey’s multiple comparisons test with α = 0.05. Outliers were identified and excluded using Grubb’s test (α = 0.05). Graphs were generated using Prism v10.3.1.

