## Supplemental for "Mouse Fc-FcγRIV structure guides Fc engineering for cross-species FcγR recognition"

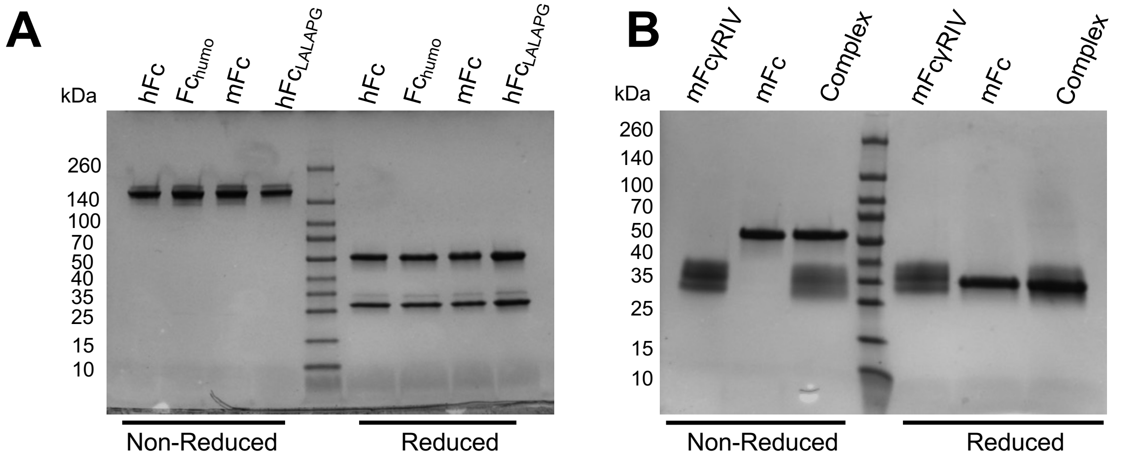

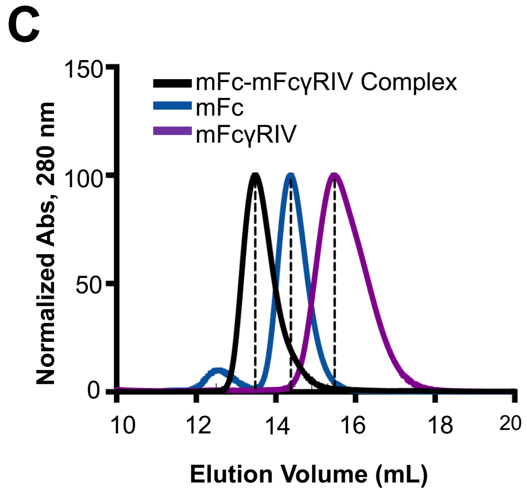

| | Elution Volume (mL) | Molecular Weight (kDa) |
|---|---|---|
| mFc-mFcγRIV Complex | 13 | 96 |
| mFc | 14 | 63 |
| mFcγRIV | 15 | 36 |

**Figure S1. Antibodies, and mouse FcγRIV, mIgG2a Fc and their complex are pure and monodisperse**.
**(*A*)** Full-length antibodies comprised of human 4D5 Fabs with mFc and hFc were analyzed by SDS-PAGE (4-20%) in reduced and non-reduced conditions.
**(*B*)** mFcγRIV, mFc (expressed without Fab arms), and their complex were similarly assessed.
**(*C*)** mFcγRIV, mFc, and the mFc-mFcγRIV complex were evaluated by SEC. Purified mFc and mFcγRIV were mixed in 1:1 equimolar ratio to obtain the complex, which was then SEC purified. The accompanying table provides peak elution volumes and their corresponding molecular weights based on SEC for each protein.

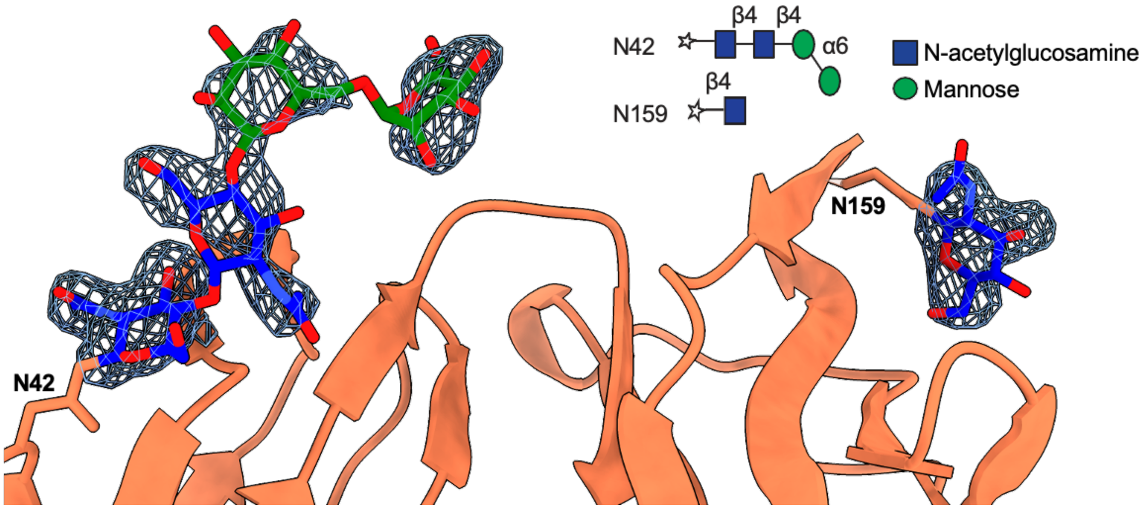

**Figure S2, related to Figure 2. Fo-Fc map validates N-linked glycan modeling in mouse FcγRIV.** 2Fo-Fc map calculated by doubling the observed electron density (Fo) and subtracting the calculated electron density (Fc) to provide an estimate of the fitting of the electron density of the N-linked glycans of mFcγRIV. The electron density is shown as blue mesh contoured to a level of 1.5 sigma (σ).

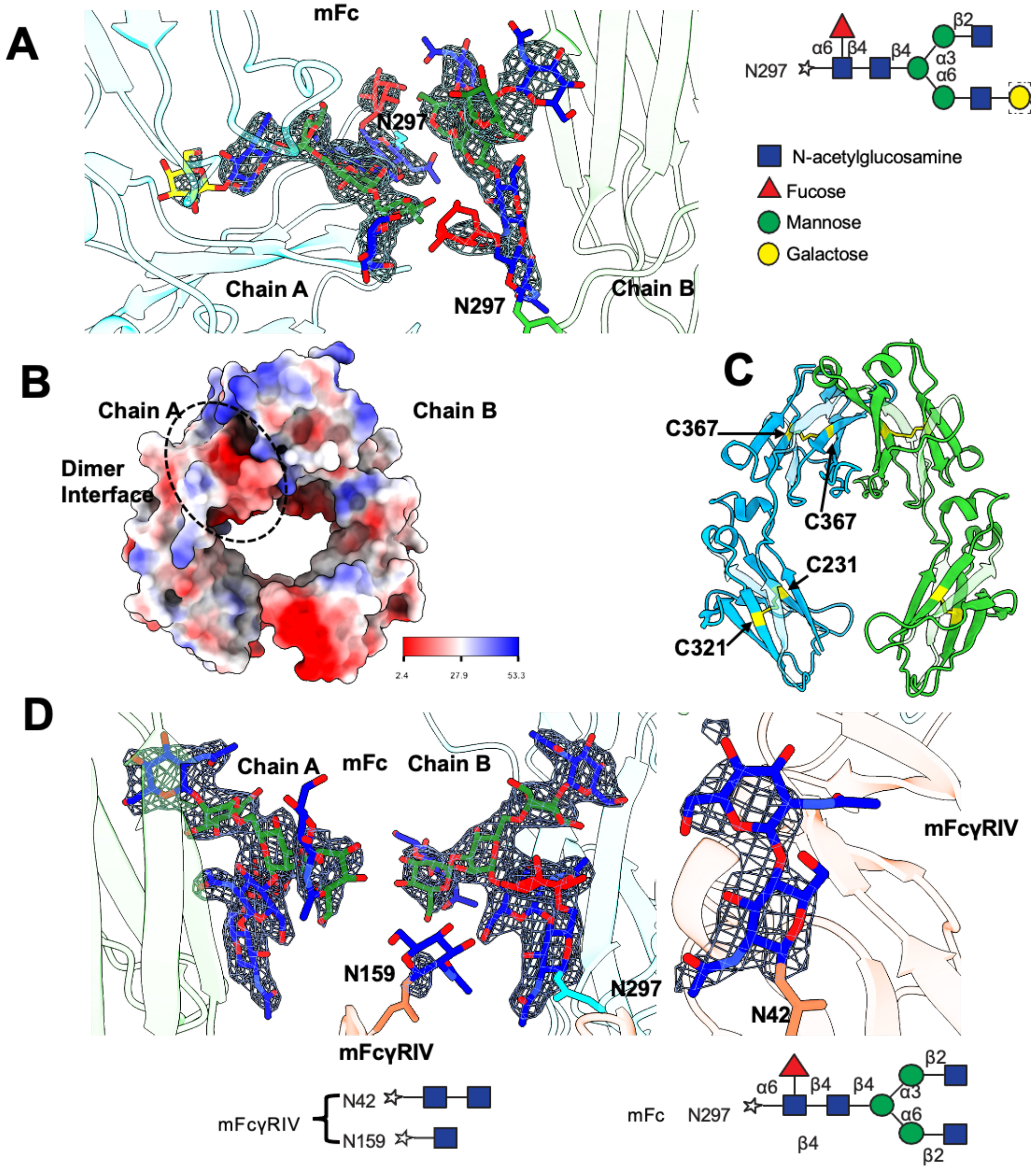

**Figure S3, related to Figure 3. Electron density and structural analyses define glycan positioning and key electrostatic features of mouse IgG2a Fc.**

(***A***) Shown here is the 2Fo-Fc map which was calculated by doubling the observed electron density (Fo) and subtracting the calculated electron density (Fc) to provide an estimate of the fitting of the electron density of the N-linked glycans of mFc. The electron density is shown as blue mesh contoured to a level of 1.5 sigma (σ).

(***B***) Electrostatic surface representation of the mFc dimer interface. The positively charged molecular surface is shown in blue, negatively charged regions in red, and neutral regions in white. A scale bar is provided to indicate the electrostatic potential.

(***C***) Standalone structure of mFc shown in the same color as Figure 2A. Residues C425, C367, C261, C321 are shown in yellow stick. Disulfide bond is shown as yellow stick.

(***D***) 2Fo-Fc map calculated by doubling the observed electron density (Fo) and subtracting the calculated electron density (Fc) to provide an estimate of the fitting of the electron density of the N-linked glycans of complex. The density is shown as blue mesh contoured to a level of 1.5 sigma (σ).

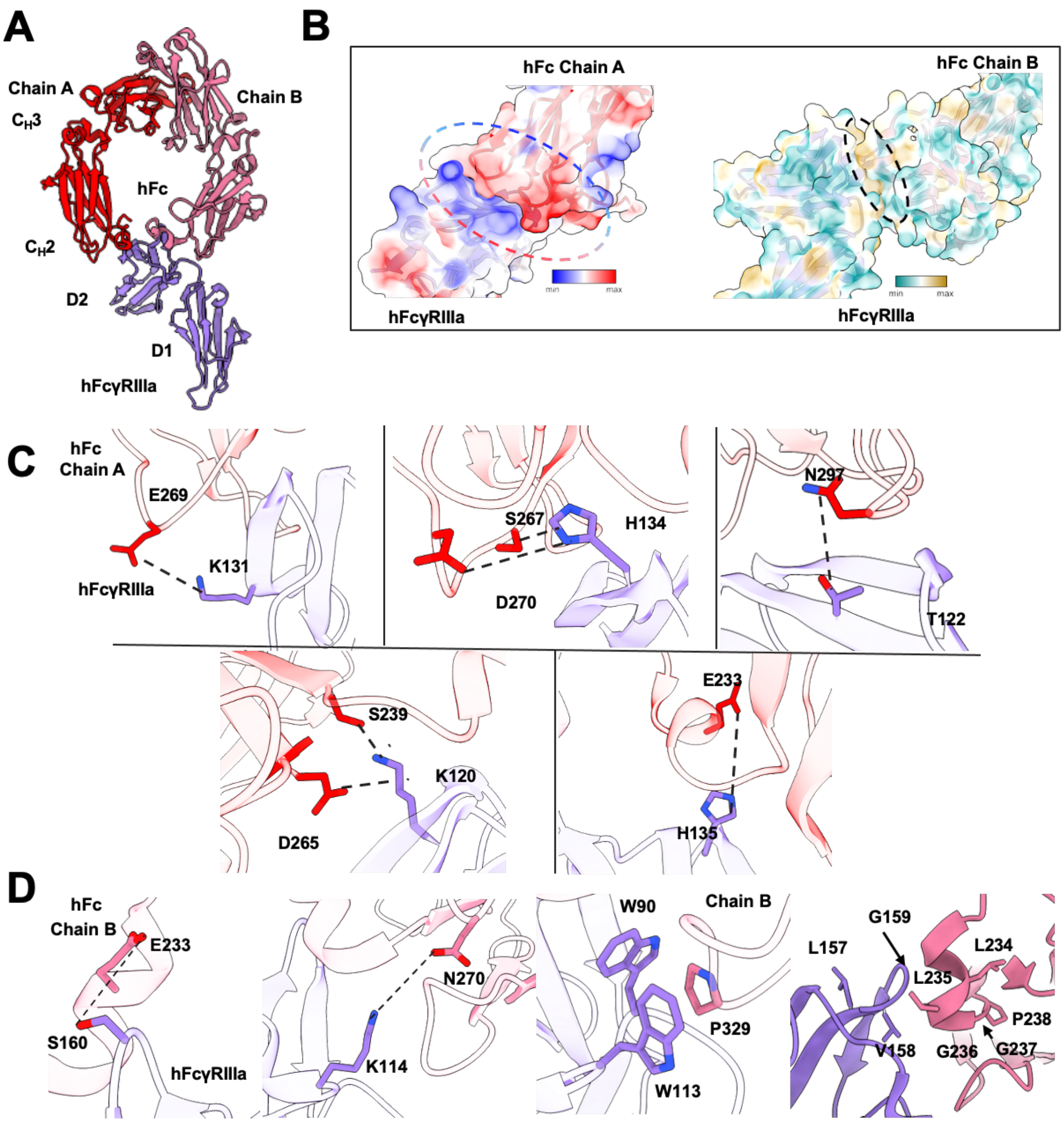

**Figure S4, related to Figure 4. Human IgG1 Fc and human FcγRIIIa complex is stabilized by important electrostatic interactions.**

(***A***) Overall structure of hFc and hFcγRIIIa (PDB ID: 3SGJ). hFcγRIIIa is shown in purple ribbon, hFc chain A is shown in red ribbon and chain B is shown in salmon ribbon.

(***B***) Interface between hFc and hFcγRIIIa, shown with electrostatic surface representation. The positively charged molecular surface is shown in blue, negatively charged regions in red, and neutral regions in white.

(***C***) Significant interactions at the interface between hFc and chain A. Key residues (E269, K131, S267, H134, D270, N297, T122, S239, D265, K120, E233 and H135) are depicted as sticks. Black dashed lines indicate salt bridges or hydrogen bonds between interacting residues.

(***D***) Significant interactions within the interface between hFc and chain B. Key residues (E233, S160, K114, N270, W90, P329, and W113) are depicted as sticks. Black dashed lines indicate salt bridges or hydrogen bonds between interacting residues.

```
                      230       240       250       260       270       280
                      |         |         |         |         |         |
hIgG1                 PAPELLGGPSVFLFPPKPKDTLMISRTPEVTCVVVDVSHEDPEVKFNWYVDGVEVHNAKT
mIgG2a                PAPNLLGGPSVFIFPPKIKDVLMISLSPIVTCVVVDVSEDDPDVQISWFVNNVEVHTAQT
hIgG1 GASDALIE        ******A**D**************************************************
hIgG1 LALAPG          ****AA******************************************************
Fchumo                ***D********************************************************
V1                    ***A********************************************************
V2                    ***N********************************************************
V3                    **************************************A*********************
V4                    **************************************E*********************
```

```
                       290       300       310       320       330       340
                       |         |         |         |         |         |
hIgG1                  KPREEQYNSTYRVVSVLTVLHQDWLNGKEYKCKVSNKALPAPIEKTISKAK
mIgG2a                 QTHREDYNSTLRVVSALPIQHQDWMSGKEFKCKVNNKDLPAPIERTISKPK
hIgG1 GASDALIE         ****************************************L*E********
hIgG1 LALAPG           ***************************************G***********
V5                     *************************************D*************
```

**Figure S5. Residue differences between human IgG1 Fc and mouse IgG2a Fc at specific positions contribute to weakened binding of human IgG1 Fc to mouse FcγRIV.** Residues in red are hFc residues involved in binding interaction with hFcγRIIIa. Underlined residues in blue are hFc residues that we hypothesized weakened interaction with mFcγRIV and were mutated to enhanced binding, resulting in hFc variants $Fc_{humo}$, V1-V5. For reference, published and experimentally verified hFc residue changes that either enhance (GASDALIE) or abolish (LALAPG) interaction with hFcγRIIIa are also shown.

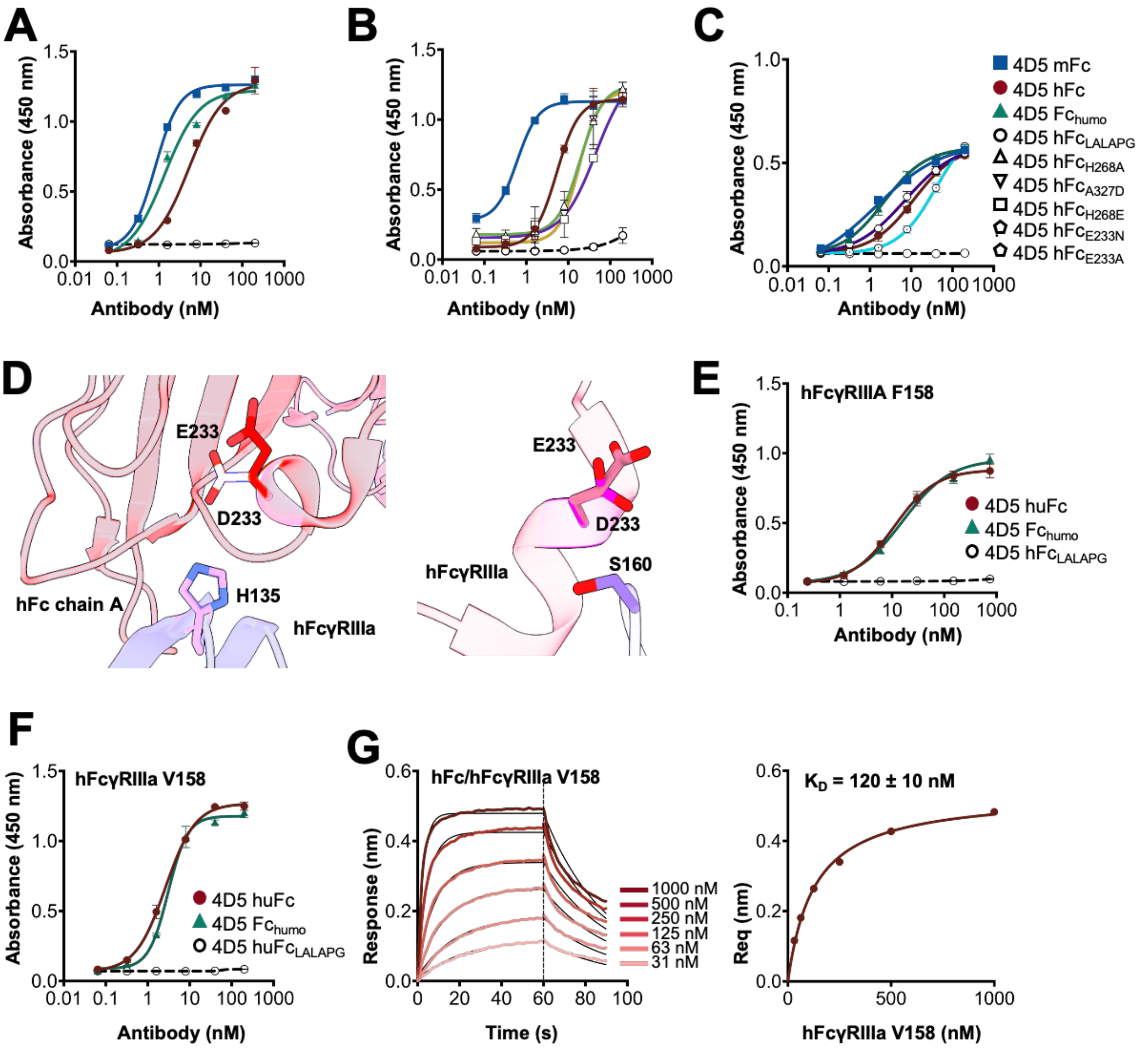

**Figure S6. Fc$_{humo}$ (E233D) mutation in human IgG1 Fc maintains binding with human FcγRIIIa and while enhancing binding to mouse FcγRIV**.

(***A-C***) ELISA data showing hFc variants with single substitutions based on structural analysis and published Fc variants to isolate a variant with enhanced binding to mFcγRIV.

(***D***) ChimeraX analyses depicting the significant interactions within the interface between hFcγRIIIa hFc chain A that are still maintained by E233D substitution.

(***E***) ELISA showing the binding of hFc and Fc$_{humo}$ with hFcγRIIIa F158 and (***F***) V158

(***G***) Biolayer interferometry (BLI) was used to quantify the binding of 4D5 hFc to hFcγRIIIa (V158) using an OctRed96 instrument. Kinetic responses for the association and disassociation phases were fit to a 1:1 binding model, while the equilibrium responses were fit to a Langmuir isotherm. $K_D$ values from kinetic and equilibrium analyses are shown in Table 1.

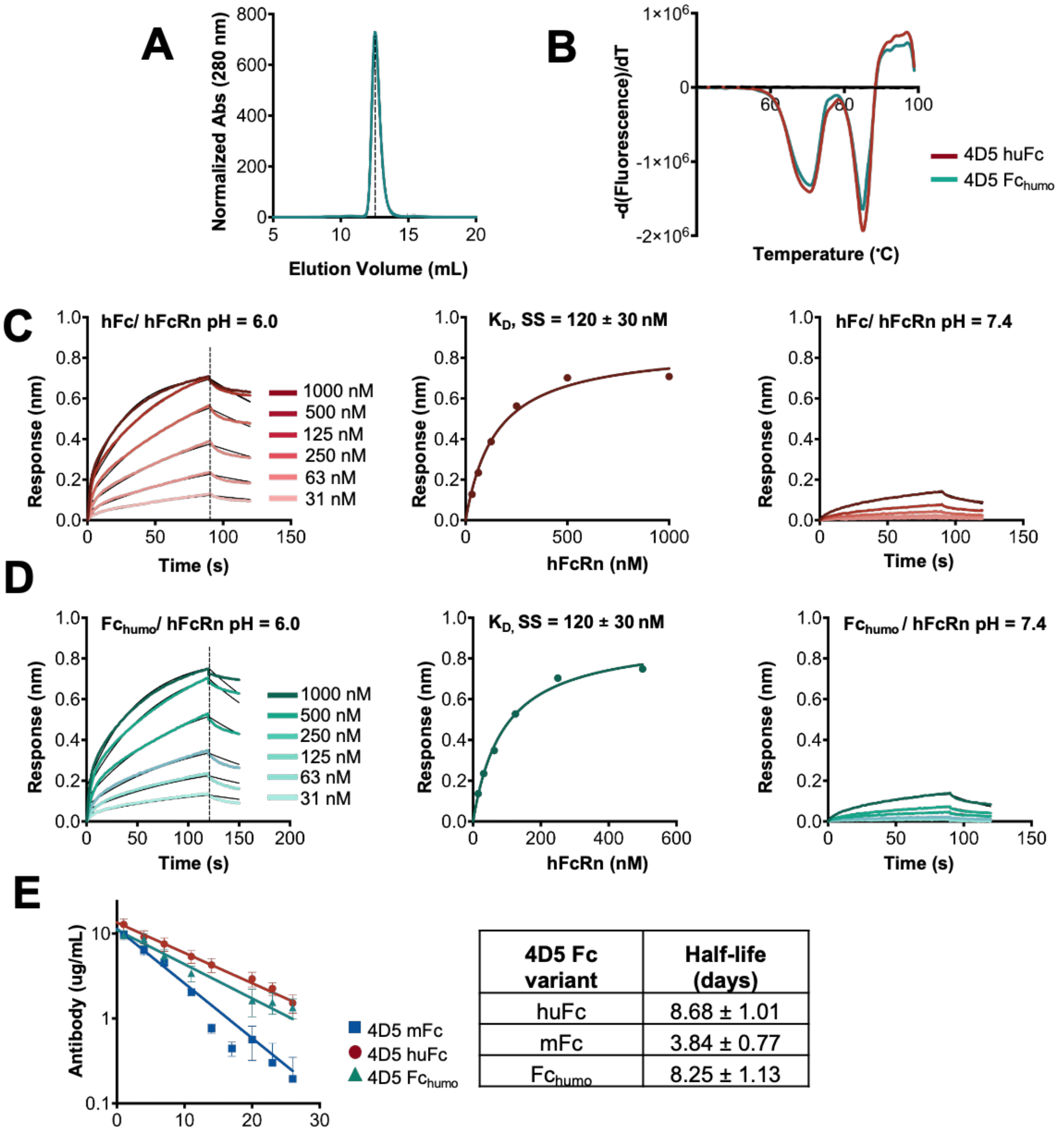

| 4D5 Fc variant | Half-life (days) |
|---|---|
| huFc | 8.68 ± 1.01 |
| mFc | 3.84 ± 0.77 |
| Fchumo | 8.25 ± 1.13 |

**Figure S7. Fc$_{humo}$ exhibits similar biophysical and pharmacokinetic properties as a human IgG1 Fc.**
**(A)** Size Exclusion Chromatography profile of 4D5 Fc$_{humo}$ shows monodispersed antibody. The elution peak at 12.54 mL corresponds to an apparent molecular weight of 149 kDa.
**(B)** Thermal stability of 4D5 hFc and Fc$_{humo}$ antibodies assessed using Protein Thermal Shift™ dye. Melt curves were measured by real-time PCR (ViiA7™) with a ramp rate of 0.05 °C/sec. Two downward peaks represent unfolding of the Fab and Fc regions, respectively. $T_m$ values for the Fc region are 70.87 °C (4D5 hFc) and 71.82 °C (4D5 Fc$_{humo}$).
**(C)** Biolayer interferometry (BLI) was used to quantify the binding of 4D5 hFc and **(D)** 4D5 Fc$_{humo}$ to hFcRn at pH 6.0 (*Left two panels*) and pH 7.4 (*Right*) using an OctRed96 instrument. Kinetic responses for the association and disassociation phases were fit to a 1:1 binding model, while the equilibrium responses were fit to a Langmuir isotherm. $K_D$ values from kinetic and equilibrium analyses are shown in Table 1.
**(E)** Serum antibody concentrations were determined by HER2-Strep ELISA and fit to determine the the β-decay rate and serum half-life of hFc, Fc$_{humo}$, and mFc. Lines were fitted with average decay for each group in GraphPad Prism. The half-life (days) is shown in the table on the right.

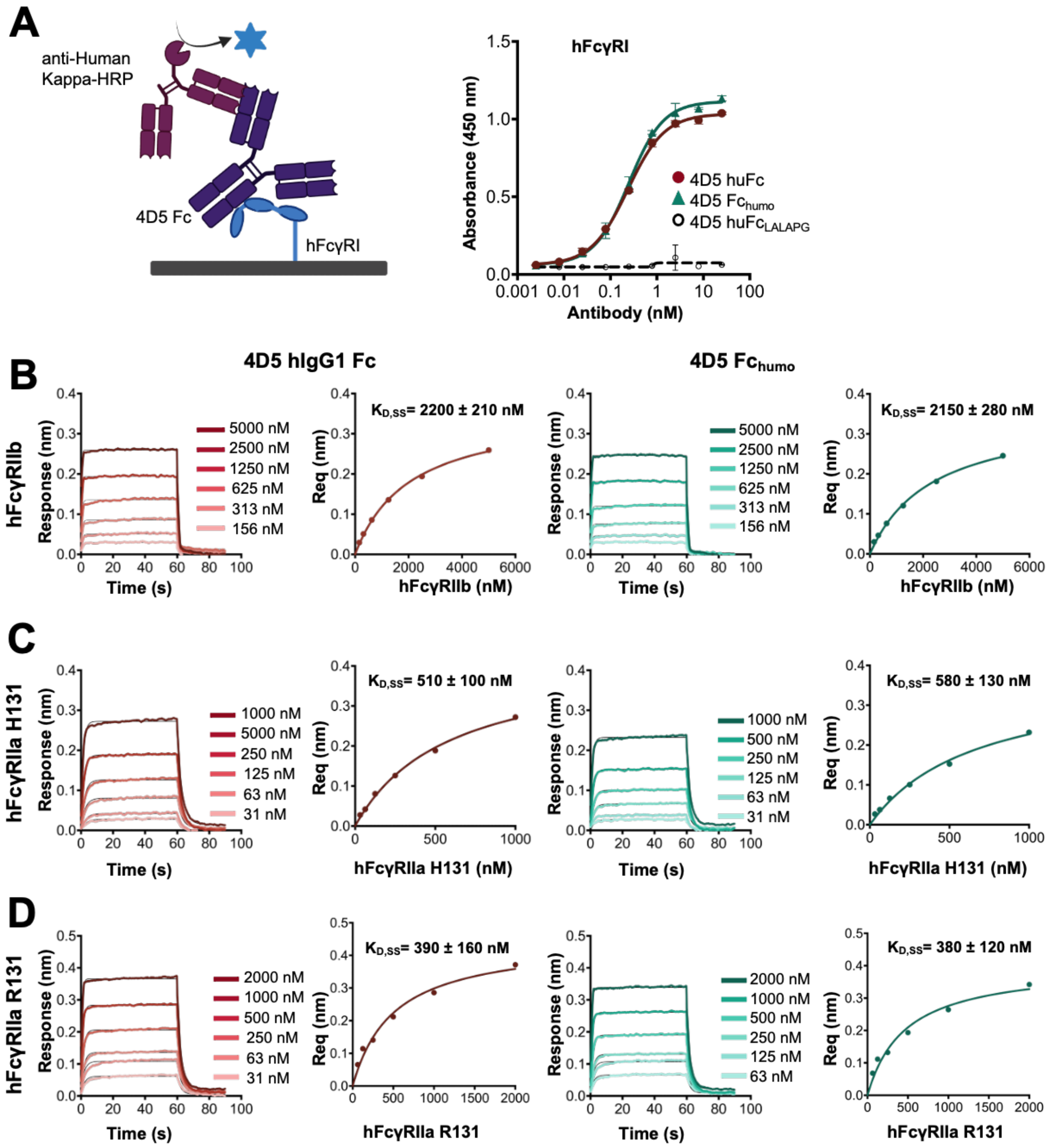

**Figure S8. $Fc_{humo}$ has similar binding to human FcγRs compared to human IgG1 Fc.**
**(*A*)** (*Left*) Schematic of indirect receptor-binding ELISA used to measure 4D5 antibodies with different Fc binding to hFcγRI. (*Right*) ELISA result showed similar binding of $Fc_{humo}$ with hFcγRI compared to hFc. **(*B-D*)** Biolayer interferometry (BLI) was used to quantify the binding of 4D5 hFc (*Left two panels*) and $Fc_{humo}$ (*Right two panels*) with (***B***) hFcγRIIb, (***C***) hFcγRIIa H131, and (***D***) hFcγRIIa R131 using an OctRed96 instrument. Kinetic responses for the association and disassociation phases were fit to a 1:1 binding model, while the equilibrium responses were fit to a Langmuir isotherm. $K_D$ values from kinetic and equilibrium analyses are shown in Table S3.

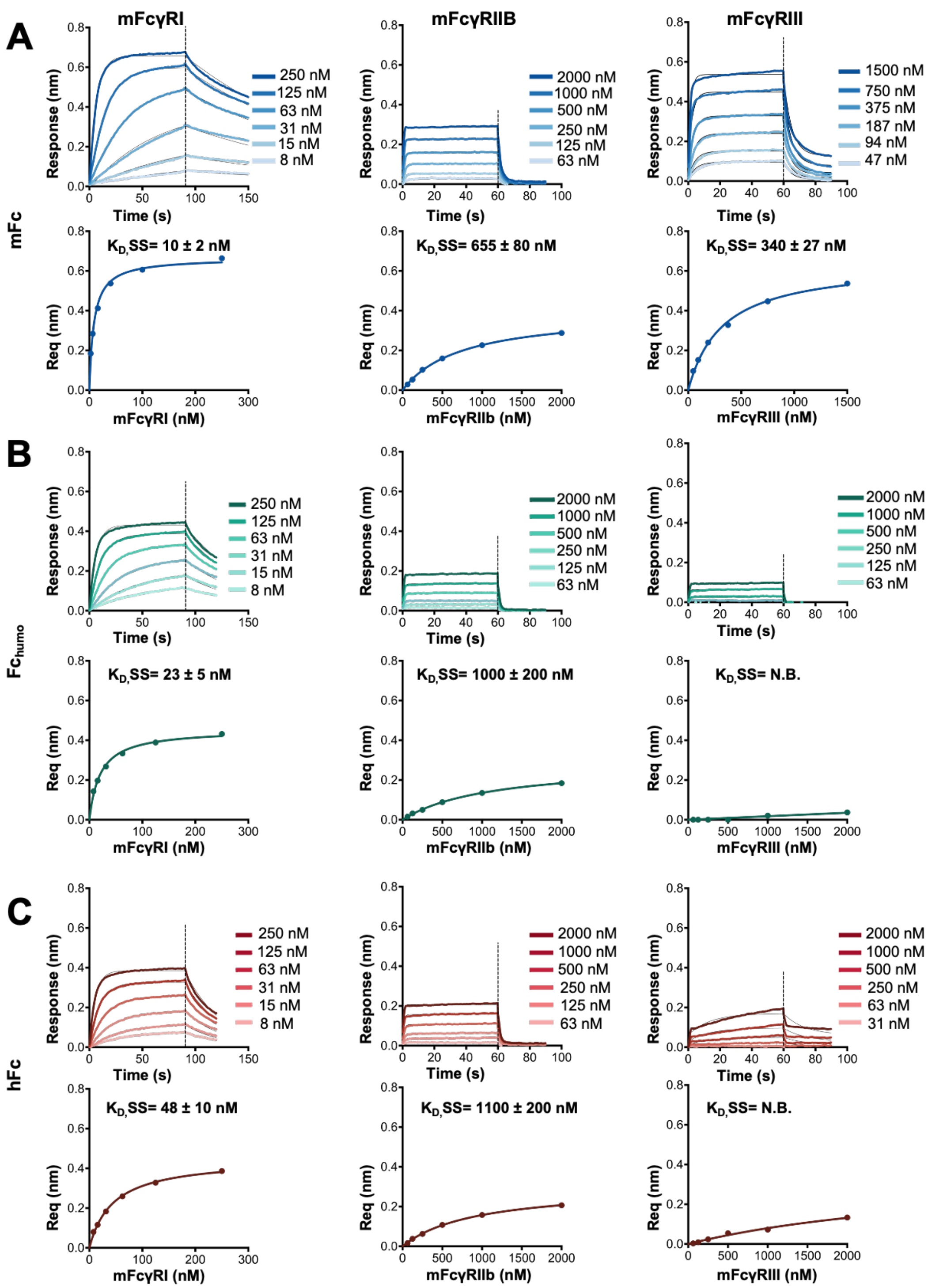

**Figure S9. Fc$_{humo}$ exhibits enhanced binding to mouse FcγRI compared to human IgG1 Fc.** (***A-C***) BLI was performed using FAB2G biosensor tips to capture (***A***) mFc, (***B***) Fc$_{humo}$, and (***C***) hFc antibody variants and then dipped into serially diluted (*Left panels*) mFcγRI (*Middle Panels*) mFcγRIIb, and (*Right Panels*) mFcγRIII at various concentrations using an OctRed96 instrument. Kinetic responses were fit to a 1:1 binding model, while the equilibrium responses were fit to a Langmuir isotherm. The data are representative of three independent replicates. The obtained $K_D$ values from kinetic and steady-state analyses are shown in Table S3. For all Fc-mFcγRIIb interactions, kinetic $K_D$ were not calculated due to a very rapid disassociation rate. No binding was determined for hFc and Fc$_{humo}$ interaction with mFcγRIII.

**Table S1: Data collection and refinement**

| | mFc | mFcγRIV | mFc- mFcγRIV Complex |
|---|---|---|---|
| **Wavelength (Å)** | 0.9999 | 0.9999 | 0.6199 |
| **Resolution range (Å)** | 66.69 - 3.21 (3.325 - 3.21) | 38.54 - 2.81 (2.911 - 2.81) | 54.23 - 3.16 (3.273 - 3.16) |
| **Space group** | C 1 2 1 | P 1 | C 2 2 $2_1$ |
| **Unit cell: a, b, c (Å)** | 135.2 98.1 61.9 | 43.01 51.17 69.08 | 113.754 185.055 93.7337 |
| **Unit cell: $\alpha$, $\beta$, $\gamma$ (°)** | 90 99.5 90 | 68.34 71.9 76.14 | 90 90 90 |
| **Total reflections** | 44101 (4319) | 21837 (2072) | 45912 (4680) |
| **Unique reflections** | 13103 (1254) | 12225 (1180) | 15713 (1611) |
| **Multiplicity** | 3.4 (3.4) | 1.8 (1.8) | 2.9 (2.9) |
| **Completeness (%)** | 99.57 (96.98) | 97.21 (93.35) | 90.48 (93.71) |
| **Mean I/sigma(I)** | 11.40 (1.56) | 4.28 (1.50) | 9.52 (2.42) |
| **Wilson B-factor(Å²)** | 81.3 | 58.5 | 53.2 |
| **R-merge** | 0.1755 (0.9197) | 0.111 (0.4289) | 0.1937 (0.6257) |
| **R-meas** | 0.2088 (1.092) | 0.157 (0.6066) | 0.2343 (0.7592) |
| **R-pim** | 0.112 (0.5843) | 0.111 (0.4289) | 0.1301 (0.4242) |
| **$CC_{1/2}$** | 0.975 (0.39) | 0.972 (0.776) | 0.938 (0.585) |
| **CC*** | 0.994 (0.749) | 0.993 (0.935) | 0.984 (0.859) |
| **Refinement** | | | |
| **Reflections used in refinement** | 13086 (1253) | 12216 (1180) | 15705 (1610) |
| **Reflections used for R-free** | 626 (50) | 612 (54) | 1552 (153) |
| **R-work** | 0.2393 (0.3255) | 0.2434 (0.3431) | 0.2263 (0.2960) |
| **R-free** | 0.2901 (0.3573) | 0.3031 (0.5529) | 0.2715 (0.3188) |
| **CC (work)** | 0.909 (0.493) | 0.835 (0.307) | 0.914 (0.721) |
| **CC (free)** | 0.884 (0.431) | 0.832 (0.193) | 0.870 (0.747) |
| **Number of non-hydrogen atoms** | 5308 | 2820 | 4897 |
| **Macromolecules** | 4989 | 2706 | 4657 |
| **Ligands** | 319 | 114 | 240 |
| **Protein residues** | 618 | 334 | 578 |
| **RMS (bonds) (Å)** | 0.014 | 0.013 | 0.012 |
| **RMS (angles) (°)** | 1.69 | 1.23 | 1.66 |
| **Ramachandran favored (%)** | 98.69 | 99.39 | 98.25 |
| **Ramachandran allowed (%)** | 1.31 | 0.61 | 1.75 |
| **Ramachandran outliers (%)** | 0.00 | 0.00 | 0.00 |
| **Rotamer outliers (%)** | 0.85 | 0.99 | 0.74 |
| **Average B-factor(Å²)** | 74.5 | 52.8 | 51.7 |
| **Macromolecules** | 73.2 | 51.8 | 51.1 |
| **Ligands** | 95.6 | 78.1 | 63.8 |
| **Molprobity score** | 16.24/97th percentile* | 10.98/97th percentile* | 13.14/95th percentile* |

*Statistics for the highest-resolution shell are shown in parentheses. $CC_{1/2}$ is the Pearson correlation coefficient for a random half of the data, the two numbers represent the lowest and highest resolution shell, respectively. MolProbity score is calculated by combining clashscore with rotamer and Ramachandran percentage and scaled based on X-ray resolution. The percentage is calculated with 100th percentile as the best and 0th percentile as the worst among structures of comparable resolution.

**Table S2. Glycan-glycan interactions in mouse and human Fc-FcγR complex**

| **hFc-hFcγRIIIa complex (PDB: 3SGJ)** | | |
|---|---|---|
| **hFcγRIIIa N162** | **hFc** | **Distance (Å)** |
| GlcNAc1 O3 | GlcNAc1 O7 (N297) | 3.3 |
| GlcNAc1 O6 | GlcNAc1 O6 (N297)<br>GlcNAc1 O5 (N297)<br>Fuc O5 (N297) | 4.2<br>4.3<br>2.9 |
| Man7 O5 | Y296 | 4.8 |
| **mFc-mFcγRIV complex** | | |
| **mFcγRIV N159** | **mFc** | **Distance (Å)** |
| GlcAc1 O3 | Y296 | 4.1 |

**Table S3. Binding constants of 4D5 Fc variants to human FcγRs.**

| | **hFcγRIIb** | | **hFcγRIIa H131** | | **hFcγRIIa R131** | |
|---|---|---|---|---|---|---|
| **Fc variants** | **$K_D$ ± SD (nM)** | **$K_{D,SS}$ ± SD (nM)** | **$K_D$ ± SD (nM)** | **$K_{D,SS}$ ± SD (nM)** | **$K_D$ ± SD (nM)** | **$K_{D,SS}$ ± SD (nM)** |
| 4D5 hFc | N.D | 2200 ± 200 | 700 ± 100 | 500 ± 100 | 600 ± 30 | 390 ± 160 |
| 4D5 $Fc_{humo}$ | N.D | 2200 ± 300 | 680 ± 140 | 580 ± 130 | 680 ± 170 | 380 ± 120 |

Data shown here are average (n=2) ± SD; N.B.= No binding detected. N.D. = $K_D$ not determined

**Table S4. Binding constants of 4D5 Fc variants to mouse FcγRs.**

| | **mFcγRI** | | **mFcγRIIb** | | **mFcγRIII** | |
|---|---|---|---|---|---|---|
| **Fc variants** | **$K_D$ ± SD (nM)** | **$K_{D,SS}$ ± SD (nM)** | **$K_D$ ± SD (nM)** | **$K_{D,SS}$ ± SD (nM)** | **$K_D$ ± SD (nM)** | **$K_{D,SS}$ ± SD (nM)** |
| 4D5 mFc | 10 ± 2 | 10 ± 2 | N.D. | 660 ± 80 | 400 ± 60 | 340 ± 30 |
| 4D5 hFc | 50 ± 8 | 50 ± 10 | N.D. | 1100 ± 200 | N.B | N.B |

Data shown here are average (n=3) ± SD; N.B.= No binding detected. N.D. = $K_D$ not determined

**Table S5. Signaling activity in biological reporter assay**

| 4D5 Fc variant | Concentration resulting in 50% max response ($EC_{50}$) (nM) | Fold induction at saturation relative to control |
|---|---|---|
| **hFcγRIIIa (V158) assay** | | |
| 4D5 hFc | 0.24 ± 0.18 | 63 ± 6 |
| 4D5 $Fc_{humo}$ | 0.29 ± 0.15 | 55 ± 11 |
| **mFcγRIV assay** | | |
| 4D5 mFc | 0.75 ± 0.35 | 22 ± 5 |
| 4D5 $Fc_{humo}$ | 1.04 ± 0.65 | 9 ± 3 |
| 4D5 hFc | N.D. | 2.1 ± 0.6 |

Data shown here are average (n=3) ± SD, with EC50 values reported with two significant figures; N.D. = $EC_{50}$ not determined due to absence of measurable dose-dependent response.
